# Transcriptomic signatures of nine high-risk neuropsychiatric copy number variants in human brain cells

**DOI:** 10.1101/2024.09.19.613680

**Authors:** Gabi Dugan, Nirmala Akula, Stefano Marenco, Pavan K Auluck, Armin Raznahan, Siyuan Liu, Alex DeCasien, Qing Xu, Ningping Feng, Bhaskar Kolachana, Xueying Jiang, Michael D Gregory, Karen F Berman, Gabriel Hoffman, Panos Roussos, Francis J McMahon, Anton Schulmann

**Author notes:** **Lead Contact**: Gabi Dugan,. Denotes equal contribution (Francis J McMahon, Anton Schulmann).

## Abstract

Large, recurrent copy number variants (CNVs) are among the strongest risk factors for neuropsychiatric conditions, contributing to multiple phenotypes with overlapping psychiatric and cognitive symptoms. However, the molecular basis of this convergent risk remains unknown. We evaluated the human brain transcriptome in carriers of nine high-risk neuropsychiatric CNVs and matched non-carriers using single nucleus RNA-sequencing. Brain tissue from carriers displayed widespread disruptions of gene expression, with thousands of differentially expressed genes, mostly located outside of the respective CNV regions. There were greater changes in deletions compared to reciprocal duplications. Functional enrichment analysis revealed changes in mitochondrial energy metabolism and synaptic function that converged across CNVs and cell types. For mirror CNVs, the direction of effects was often reversed between deletions and duplications and showed correlation with CNV gene dosage. These findings suggest that a shared pathophysiology underlies risk for convergent brain phenotypes across CNVs and point toward promising therapeutic targets.

## Introduction

Copy number variants (CNVs) are chromosomal alterations consisting of a deletion or duplication of specific segments on the chromosome (1,2). This type of variant is widespread across the genome and a significant source of functional genetic variation (1,3,4). CNVs can range in size from a few kilobases to several megabases, and are thought to raise disease risk through changes in gene dosage (5). Some large, rare but recurrent CNVs are associated with substantial risk for various neuropsychiatric conditions, such as autism spectrum disorder, intellectual disability, schizophrenia, and mood disorders (6–13), and carriers of these neuropsychiatric CNVs exhibit symptoms in the domains of cognition, perception, and mood. Overlapping diagnoses and symptoms among carriers of distinct CNVs suggest some shared pathophysiology (9,14,15), but the molecular basis of this convergent risk remains largely undiscovered.

Highly penetrant CNVs offer a unique opportunity to explore the impact of dosage changes in specific sets of genes within CNV regions. While many neuropsychiatric CNVs have been extensively studied with cell culture, mouse models, and neuroimaging methods (16–19), little is known about the cellular and molecular effects of CNVs within the human brain itself. Studies in human tissue have so far been limited to bulk analysis (20,21). Through advances in single-nucleus RNA-sequencing, it is now possible to interrogate the molecular effects of CNVs in individual cell types, revealing cell type-specific changes that may be masked in bulk tissue analyses.

In this study, we used single-nucleus RNA-sequencing of post-mortem human brain from individuals harboring rare, high-risk CNVs to explore the neurobiological mechanisms through which these CNVs confer risk for neuropsychiatric illnesses. We analyzed human brain tissues from donors with pathogenic deletions and duplications on chromosomes 22q11.2, 16p11.2, 15q11.2, or 1q21.1, and deletions on 7q11.23, along with matched non-carriers, derived from three brain banks. This unique collection encompassed some of the most penetrant genetic risk factors for neuropsychiatric illnesses, including four “mirror” CNVs with reciprocal deletions and duplications in the same locus. Cell type specific gene expression changes shed light on neurobiological perturbations associated with these CNVs and reveal convergent mechanisms that may underlie overlapping patterns of psychopathology.

## Methods

### Human brain tissue samples

Human brain samples were derived from three brain banks: the NIMH Human Brain Collection Core (HBCC), the University of Maryland Brain and Tissue Bank (UMB), and the Stanley Medical Research Institute Brain Collection (SMRI). All brain samples were obtained using protocols approved by the Institutional Review Boards (IRB) with permission of the next-of-kin. Psychiatric diagnoses were inferred based on next-of-kin interviews and medical records, whenever available, according to DSM-IV and DSM-V criteria.

### Genotyping and CNV Calling

Genotyping was performed as previously described (79) using a combination of Human1M-Duo (v3), HumanHap650Y (v3), HumanOmni5-Quad and Global Screening Array BeadChips. To call CNVs, we used PennCNV (80) with standard settings. We examined if any of the CNV calls overlapped with known CNVs previously associated with neuropsychiatric disorders (7). B-allele frequencies and log R ratios were then plotted for the respective CNV gene to confirm CNV status (Figure S1). (For more information on CNV carrier selection and matching approaches see Supplementary Methods) Since the exact CNV breakpoints could not be precisely determined, the ClinGen(24) coordinates of the corresponding CNVs were used (Table S1, 2). Quality-related variables across the dataset are summarized in Table S4.

### Nuclei isolation

50 mg of frozen brain tissue were dissociated using Singulator 100 (S2 Genomics) in a prechilled nuclei isolation cartridge, following the Extended Nuclei Isolation protocol provided by the manufacturer. Nuclei were collected by centrifugation at 500g for 5 min at 4°C in a 15 mL tube, washed with 3 mL nuclei storage reagent (NSR) once, then washed with 3 mL washing buffer (PBS + 1% BSA + 0.2 U/μL RNase inhibitor) and filtered through 30 μm MARC SmartStrainer (Miltenyi Biotec) twice, resuspend in 800 μL washing buffer and filtered through 40 μm Flowmi cell strainer (Bel-Art) before loading.

### Nuclei capture and library construction

Nuclei were captured and single-nucleus RNA-sequencing libraries were prepared following the 10X Genomics protocol CG000315 RevC: Chromium Next GEM Single Cell 3’ Reagent Kits v3.1 (Dual Index). After washing and nuclei counting, approximately 8,000 nuclei from each sample were loaded on the 10X Genomics Chromium Controller. The Reverse Transcription and cDNA amplification were done following the manufacturer’s protocol. 3’ mRNA-seq gene expression libraries were prepared from 100ng cDNA on Sciclone G3 Liquid Handling Workstation (Revivity).

### Next Generation Sequencing

Libraries were pooled and sequenced on NovaSeq S4 100 cycle flow cell with 2% PhiX spiked in. The paired-end dual index sequencing run was set up as Read1 (28 cycles, i7 Index: 10 cycles, i5 Index: 10 cycles) and Read 2 (90 cycles).

### Single-Nucleus RNA-Sequencing Data Processing

Using the 10X Genomics CellRanger’s processing software (v 7.0.0) (25), we demultiplexed the raw base call (BCL) files. The generated FASTQ files from CellRanger were aligned to the reference human genome GRCh38p14 to create raw feature-barcode count matrices. Quality control, clustering, and cell type annotations are detailed in Supplementary Methods.

### Differential Gene Expression

Differential gene expression (DGE) analysis was achieved using dreamlet (v 0.99.6) following default metrics (26) (see *Supplementary Methods* for further description).

To explore if or which other covariates should be included in the DGE model, we used variance partitioning (27) and linear regression analyses where we iteratively added in covariates.

### We chose the following DGE model in dreamlet

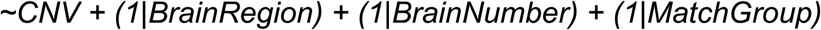

We also removed highly abundant mitochondrial-encoded and ribosomal genes that are often associated with sample quality: gene names starting with “MT-”, “RPL”, and “RPS”.

While no covariate was significantly associated (p-value <0.05) with PC1 for all nine cell types, we did run a quality-covariate adjusted model to ensure the findings did not change (see *Supplementary Methods*).

To test for CNV gene dosage effects, we split carriers into groups by CNV, comprised of deletions, reciprocal duplications, and kept all non-carriers. We then classified samples by CNV gene dosage, with deletions classified with a 1, non-carriers a 2, and duplications a 3. This continuous gene dosage classification was then used as the fixed effect in the model, keeping the same grouping factors:

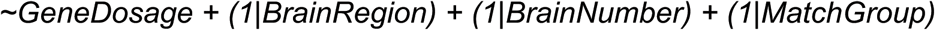

This yielded genes that were positively associated with CNV gene dosage (positive logFC) and negatively associated with CNV gene dosage (negative logFC).

### Functional Enrichments

Rank-based functional enrichment testing approaches were used to mitigate noise and unreliable results due to arbitrary p-value thresholds, known to influence threshold-based functional enrichment testing results (28,29). GSEA was our primary functional enrichment testing method as it is widely used and acknowledged to be a strong candidate for discovering biological insights for RNA-seq data analysis (28,30). The GSEA algorithm was implemented using fgsea (v 1.24) (31).

We tested for functional enrichments using Gene Ontology (GO) Biological Processes (BP) database (32–35). Mitocarta (36) was used to obtain more detailed curated terms related to mitochondrial functions, including OXPHOS complexes. We also used SynGO to select terms related to the synapse (37).

Results were adjusted for multiple testing using the Benjamini-Hochberg procedure.

### Infant Samples Clustering, Cell Type Annotations

We also analyzed a smaller infant dataset, which included samples from the dlPFC and ACC of infant carriers of the 7q11.23 (WBS) or 15q11.2 (BP1-BP2) deletion, along with matched infant non-carriers (Figure 1A). QC steps and analyses are described in the supplementary material.

**Figure 1:**
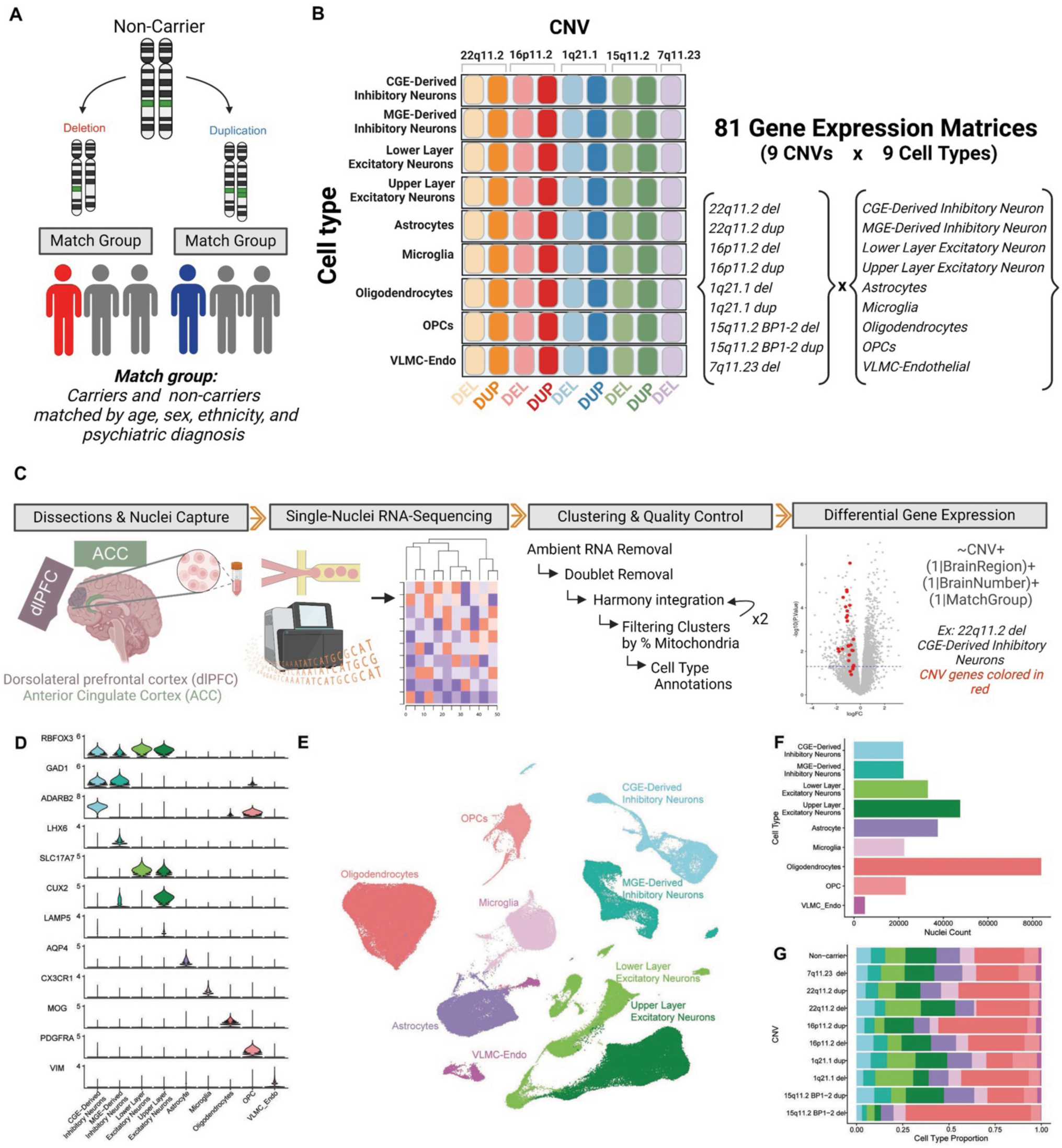
Overview of single-nucleus transcriptomic workflow we used to explore the effects of high-risk CNVs in human brain cells. (A) Outline of study design. Each of 15 carriers (13 adults and 2 infants) across nine CNVs, was matched with two non-carrier individuals (match groups). (B) Infographic of dataset, which comprised 9 CNVs and 9 cell types, leading to 81 differential gene expression (DGE) tests. (C) Overview of data analysis pipeline. Dorsolateral prefrontal cortex (dlPFC) and anterior cingulate cortex (ACC) were dissected, nuclei were captured and underwent single-nucleus RNA-sequencing. Sequenced data was analyzed with iterative clustering and quality control. Resolved clusters were annotated into nine well-resolved cell types using Azimuth (90,91) . Differential gene expression analysis was conducted between carriers and non-carriers accounting for brain region, brain sample number, and match group as random effects using dreamlet (26). (D) Stacked violin plot showing normalized expression of selected marker genes for each major cell type. (E) UMAP visualization of single nuclei colored by annotated cell types. (F) Barplot depicting total number of nuclei for each of the nine cell types tested (G) Barplot depicting cell-type proportions across CNV carrier and non-carrier groups. [Created with <u>Biorender.com</u>].

## Results

### Single-Nucleus RNA-Sequencing Pipeline

We performed single-nucleus RNA-sequencing of post-mortem human brain tissue from 15 individuals (13 adults, 2 infants) carrying high-risk neuropsychiatric CNVs along with matched non-carriers (Figure S1, See Methods & Supplementary Methods). Carriers harbored deletions or duplications on chromosomes 22q11.2, 16p11.2, 15q11.2, 1q21.1, or the deletion on the 7q11.23. Each carrier was matched with two non-carriers based on brain bank, age, sex, race, and psychiatric diagnosis, resulting in a total of 28 (24 adult and four infant) non-carrier control individuals. Whenever possible, tissue from dorsolateral prefrontal cortex (dlPFC) and anterior cingulate cortex (ACC) was collected for sequencing (Figure 1A, C).

Starting with the adult samples, strict, iterative quality control was applied to remove poor quality nuclei and samples (Methods, Figure S2,3). After clustering the remaining 295,299 nuclei, we resolved nine cell types for downstream analysis: CGE- and MGE-derived inhibitory neurons, upper- and lower-layer excitatory neurons, astrocytes, microglia, oligodendrocytes, oligodendrocyte precursor cells (OPCs), and vascular leptomeningeal cells (VLMC)-endothelial cells (Figure 1C,D; Figure S4A-C, Figure S5C).The distribution of cell types was consistent across brain regions, aligning with the high degree of concordance expected for the dlPFC and ACC in broad cell types (38) (Figure S5B). The infant samples underwent a similar pipeline but were processed separately due to differences in cell type composition (Methods; Figure S6A-D). After filtering, the dataset yielded a total of 66 adult and 10 infant samples.

For differential gene expression (DGE), we used dreamlet (Version 0.99) (26). Our DGE model included match group, brain region, and subject as random effects and CNV as a fixed effect. “Match group” represents the grouping of carriers and matched non-carriers and was correlated with several other demographic variables (Figure S7A, See Methods & Supplementary Methods). An alternative model accounting for quality-related variables, namely RNA integrity number (RIN), gene detection rate, and cell type count, yielded similar results (Figure S7D, E).

### DGE model captures expected changes in CNV regions

As internal validation, we first examined DGE within each CNV region to test if gene expression changes reflected CNV gene dosage. We extracted all genes within each CNV region (Methods; Table S1,2) and evaluated the DGE statistics from each DGE test. Mean log fold-change (logFC) across cell types confirmed reduced expression of most genes within deletions and increased expression of most genes within duplications (Figure 2A; Figure S9). The expected direction is also shown in each cell type, thus capturing the anticipated direction of gene dosage change (Figure 2B; Figure S8).

**Figure 2:**
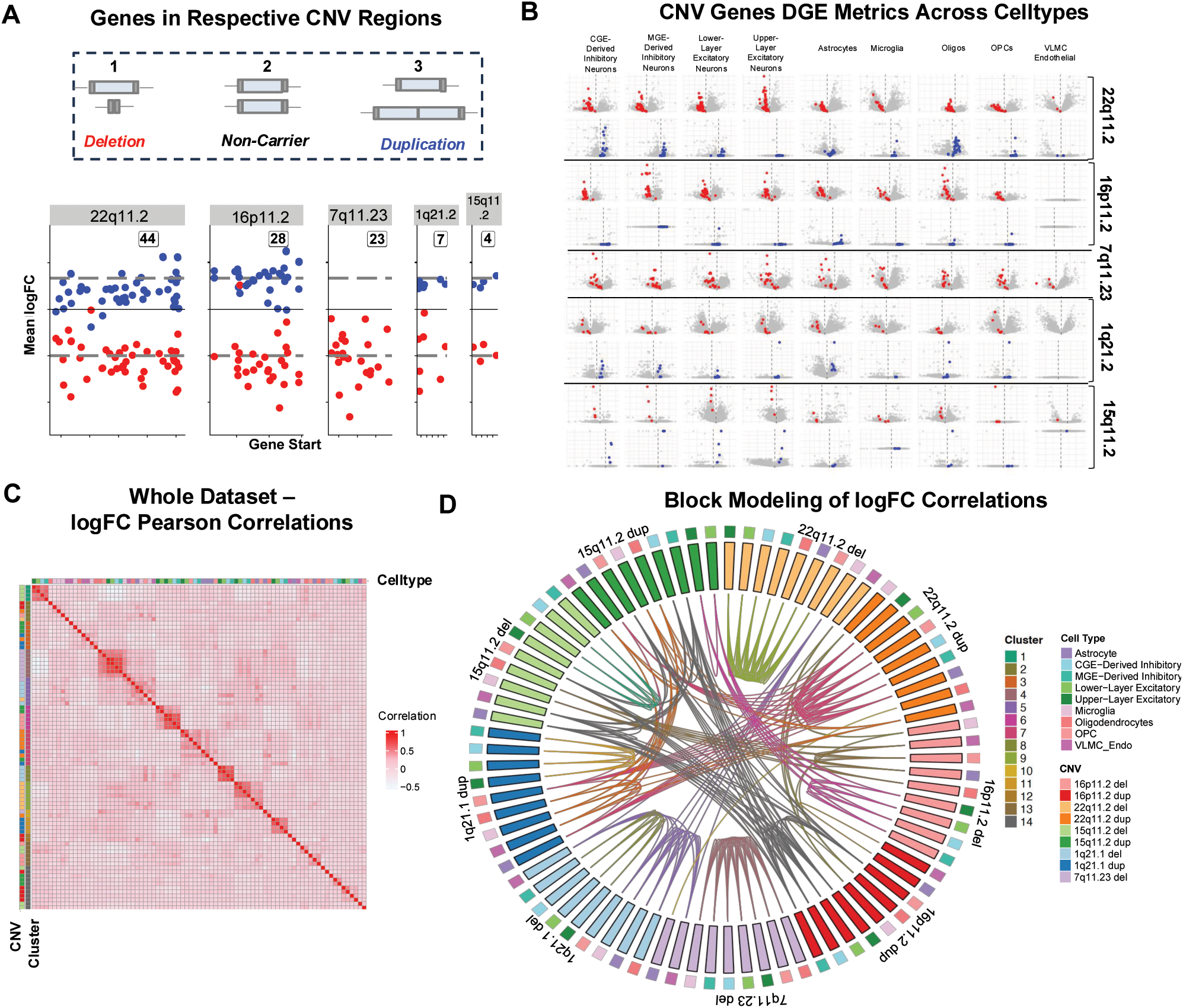
Differential expression of genes within each CNV region and across the transcriptome. CNV regions were based on ClinGen (24) in GRCh38. (A) Mean logFC (y-axis) of genes in respective CNV regions across nine cell types shown on the y-axis and respective gene positions are shown on the x-axis. Results from deletions (red) and duplications (blue) show that with few exceptions genes within deletions are downregulated (negative logFC) and genes within duplications are upregulated (positive logFC). Dotted lines indicate expected logFC based on copy number. (B) Volcano plots of DGE testing for each CNV (row) and celltype (column). CNV genes are highlighted, maintaining the deletions in red and duplications in blue color scale. (C) Heatmap showing Pearson correlations of logFC in gene expression between all pairs of CNVs across nine cell types. Each row and column represent a CNV-cell type pair, with clustering revealing blocks of similar transcriptomic effects across CNVs and cell types. (D) Circos plot showing groupings from weighted stochastic block modeling (WSBM) of the CNV-cell type correlation matrix. This highlights distinct modules of similar transcriptional impact, indicating convergent patterns of gene regulation across CNVs and cell types. Clusters are annotated by CNV (inner ring) and cell type (outer ring).’

Previous work suggested that large CNVs can affect expression of nearby genes, on the same chromosome arm (39). To explore this question in our data set, we examined the DGE t-statistics of genes in proximity to each CNV region: those on the same chromosome, on the same chromosome arm, within 5 megabases (Mb) of the CNV, and within 1 Mb of the CNV. We observed some evidence of disrupted gene expression near CNV regions, particularly in duplications (Figure S9). However, most of the DEGs lay far outside of the CNV region and surrounding areas. Thus, DGE results captured widespread effects beyond the CNV and surrounding genomic regions.

### Transcriptomic signatures show convergence across CNVs

Expanding the analysis to the whole transcriptome, we calculated the Pearson correlations of the logFC vectors across the 81 dimensions (9 CNVs × 9 cell types) from DGE (Figure 2C; Table S8). We then performed weighted stochastic block modeling (WSBM) on the matrix of Pearson correlations. WSBM identified structured communities of CNV-celltype combinations that exhibited similar transcriptomic profiles, suggesting convergence across distinct CNVs in specific cell types (Figure 2D). Notably, CNVs clustered by both CNV identity and cell class, with some glial populations forming another tightly connected module that spanned duplications and deletions. For example, glial cells from 16p11.2 and 22q11.2 deletion and duplication CNVs and from 15q11.2 dup and 1q21.1 dup CNVs showing intercorrelations, suggesting common downstream effects of distinct CNVs in non-neuronal lineages. These patterns of correlation converging across CNVs and cell types indicate that the transcriptional impact of high-risk CNVs is not uniformly idiosyncratic. Instead, specific combinations of CNVs and cell types exhibit shared gene expression changes, suggesting a potential for convergent mechanisms of neurodevelopmental risk.

### Deletions have greater impact on the transcriptome than reciprocal duplications

Having confirmed that the DGE model captured expected differences in gene copy number, we proceeded to investigate changes across the entire transcriptome (Table S8). DGE analysis revealed thousands of differentially expressed genes (DEGs) spanning cell types and CNVs. Deletions consistently yielded more differentially expressed genes (DEGs) than their reciprocal duplications. The largest DEG counts were observed in 22q11.2 (3111), 7q11.23 (2220), 16p11.2 (457), and 1q21.1 (7746) deletions, while duplications at 15q11.2, 1q21.1, 22q11.2, and 16p11.2 showed markedly fewer DEGs (9, 3, 22, and 16, respectively; Figure 3A). We found that the 22q11.2, 1q21.1, and 16p11.2 deletions all showed greater transcriptomic impacts than their reciprocal duplications (Figure 3B). A greater impact of deletions aligns with the higher penetrance and more extensive brain structural changes typically reported in deletions compared to duplications (9,40–43).

**Figure 3:**
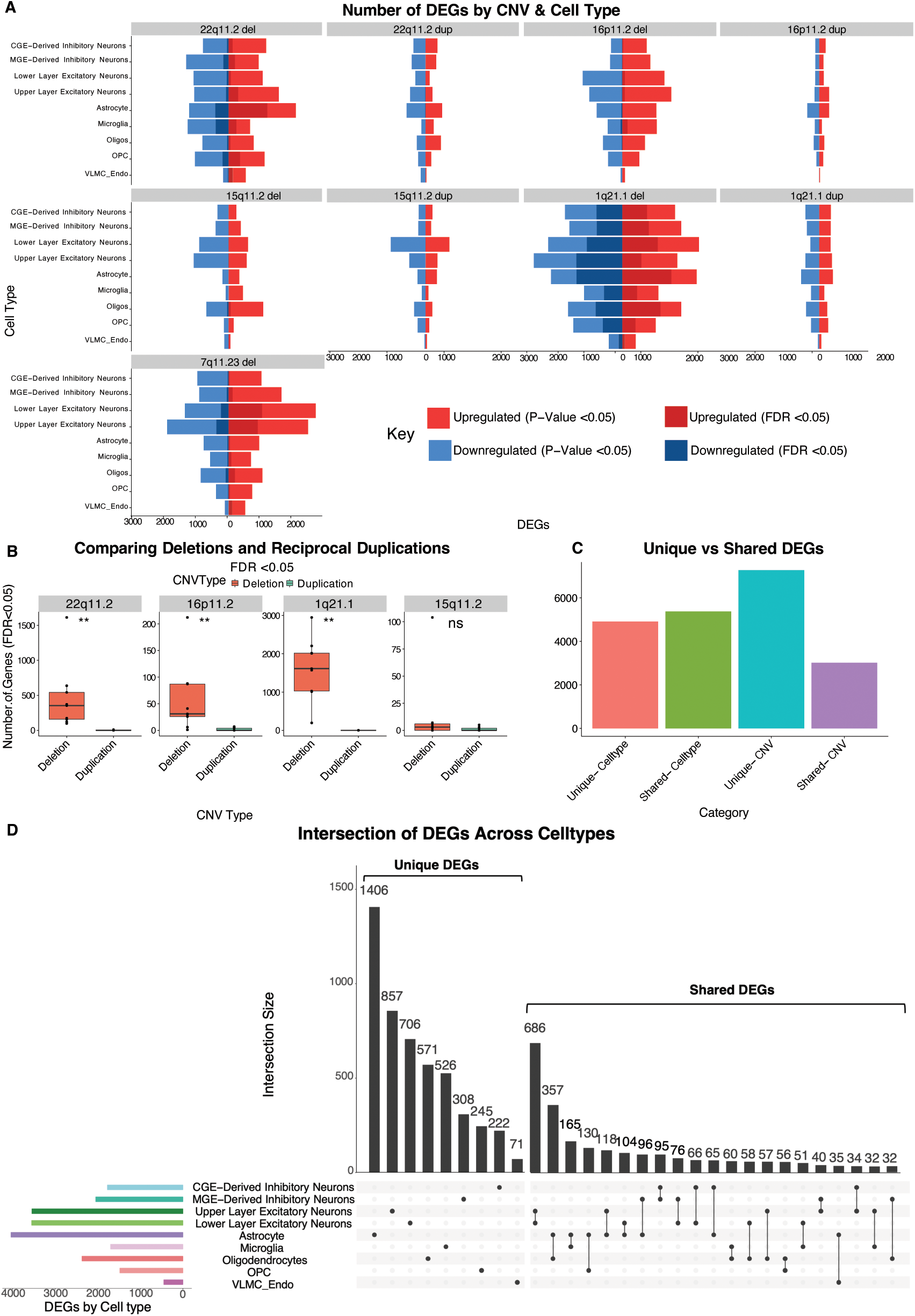
Number of differentially expressed genes (DEGs) across CNVs and cell types. (A) Number of DEGs for each CNV in all nine cell types, at p-value<0.05 (nominal) and FDR <0.05. Upregulated DEGs are shown as positive bars and downregulated DEGs as negative bars, with color indicating directionality and significance level. (B) Number of DEGs for each deletion and duplication across the 9 cell type models. Paired Mann-Whitney U-test on genes with FDR <0.05 show significant differences between deletions and reciprocal duplications. (C) Barplot showing the breakdown of differentially expressed genes (DEGs) based on whether they are unique to a single cell type or shared across multiple cell types, followed by a breakdown based on whether these DEGs are unique to a single CNV or shared across multiple CNVs. (D) Intersection of DEGs across cell types shown by Upset plot. Bars show the number of unique DEGs for each cell type first and the magnitude of the intersection of DEGs between cell type pairs second, ordered by intersection size. The top 30 intersections are shown. Bar plot on bottom left shows the union of all DEGs by cell type.

Our single-nucleus RNA-seq approach allowed us to delineate the impact of CNVs on distinct cell types. Notably, we observed the greatest differences across cell types in the 7q11.23 and 22q11.2 deletions. In the 22q11.2 deletion, astrocytes exhibited 16.4 times more DEGs than the least affected cell type, oligodendrocytes, and 2.5 times more DEGs than the second most affected celltype, microglia. In the 7q11.23 deletion, upper- and lower-layer excitatory neurons showed between 4.4 to 22.4 times more DEGs than other cell types, with OPCs showing the fewest and oligodendrocytes showing the next highest DEG burden. These data highlight astrocytes and excitatory neurons as particularly vulnerable in these high-risk CNVs, pointing to cell type-specific mechanisms of transcriptional dysregulation (Table S7).

There were 4912 genes differentially expressed (FDR <0.05) unique to one celltype, and 5378 genes shared between two or more celltypes, demonstrating that both cell-type specific and convergent gene expression changes contributed to the CNV signals (Figure 3C). Astrocytes showed the greatest number of overall (n=4038) and unique DEGs (n= 1406), and shared DEGs with other glial populations, such as oligodendrocytes, microglia, and OPCs (Figure 3D). Upper- and lower-layer excitatory neurons showed the second greatest numbers of DEGs and shared the greatest overlap with greater than 600 DEGs shared between the two excitatory neuron cell types. The large number of cell type-specific DEGs, as well as interesting overlaps, highlights the relevance of using single-cell genomics to elucidate the complex impact of these high-risk variants.

### Disruption of the transcriptome correlates with deletion size

To quantify relationships between genes within each CNV region and gene expression in the rest of the transcriptome, we scored CNVs for gene content and loss-of-function intolerance. Deletions showed a correlation between the number of genes in the CNV region and the number of DEGs (Figure S10A), whereas duplications did not (Figure S10B). We excluded the 1q21.1 CNV from this analysis due to variable involvement of the proximal TAR region (Table S1), leading to an inflated number of DEGs in the 1q21.1 deletion carriers we studied.

Gene dosage intolerance is an important predictor of pathogenic CNVs. To investigate this, we utilized the loss-of-function observed over expected upper bound fraction (LOEUF) metric, which measures tolerance to loss-of-function mutations (44–46). For each DGE model, we summed the inverse-LOEUF scores of all genes in the CNV region retained in DGE-testing (See methods). We then correlated this sum with the number of DEGs. Among deletions (again excluding 1q21.1), we observed a positive correlation between the number of DEGs and the summed inverse LOEUF scores (Figure S10C). In contrast, no correlation was detected among duplications. Thus, deletions with greater aggregated constraint metrics are associated with larger transcriptomic disruptions.

### Functional enrichments reveal convergence on mitochondrial energetics and synaptic terms across CNVs

A key objective of our study was to assess the functional impact of changes in gene dosage across CNVs and cell types and to explore functional convergences. To accomplish this, we performed rank-based functional enrichment testing for each of the 81 DGE models (9 CNVs by 9 cell types) (Table S9). To interrogate convergent signals, we evaluated significant gene ontology (GO) terms in three ways: across all 81 models (Figure 4B), for each CNV across cell types (Figure S11A), and for each cell type across CNVs (Figure 5A). We defined functional convergence here as sharing the same GO terms in functional enrichment analysis. This analysis was done irrespective of direction.

**Figure 4:**
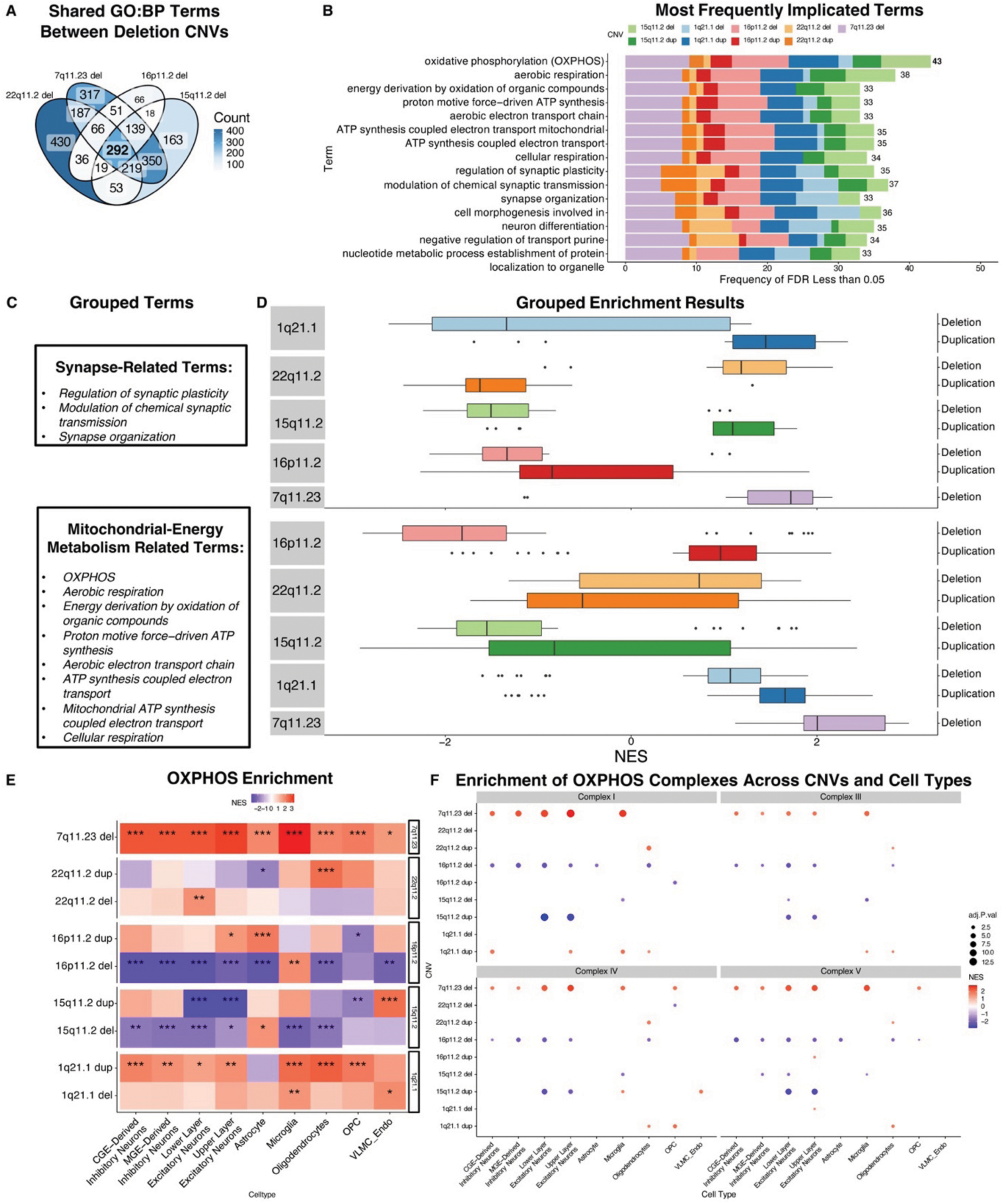
Convergence of functional enrichments across CNVs on synapse and mitochondrial energy metabolism terms. (A) Venn diagram showing the overlap of enriched Gene Ontology Biological Process (GO:BP) terms across four deletion CNVs: 15q11.2, 22q11.2, 16p11.2, and 7q11.23. (B) Barplot showing the intersection of the top fifteen most frequently implicated terms (y-axis) across the 81 tests (nine cell types and nine CNVs). (C) Groupings of the GOBP terms by broad themes. Eleven of the fifteen ten terms were related to synapses or to mitochondrial energy metabolism. (D) Boxplot depicting normalized enrichment scores (NES), which normalizes for number of genes in the term(x-axis) of synapse-related terms (aggregating three GO terms) and those related to mitochondrial energy metabolism (aggregating five GO terms). NES values are arranged on the x-axis and colored by CNV. There were observable CNV gene dosage effects with several deletions and reciprocal duplications having opposite directions of enrichments, e.g. synapse related terms are positive in 22q11.2 deletion and negative in 22q11.2 duplication. (E) Heatmap showing the OXPHOS (GO:0006119) functional enrichment results, which was the most frequently term across the nine cell types (x-axis) and nine CNVs (y-axis). The grid is colored by NES and annotated by FDR significance thresholds (FDR <0.05 = *, FDR <0.01 = **, FDR <0.001 = ***). OXPHOS is significant in all CNVs, often across cell types. (F) Enrichment of OXPHOS complexes from Mitocarta (36). Only enrichments with FDR <0.05 are shown. Complex II consists of only four genes and therefore is of limited power for enrichment testing, so was excluded from the analysis.

**Figure 5:**
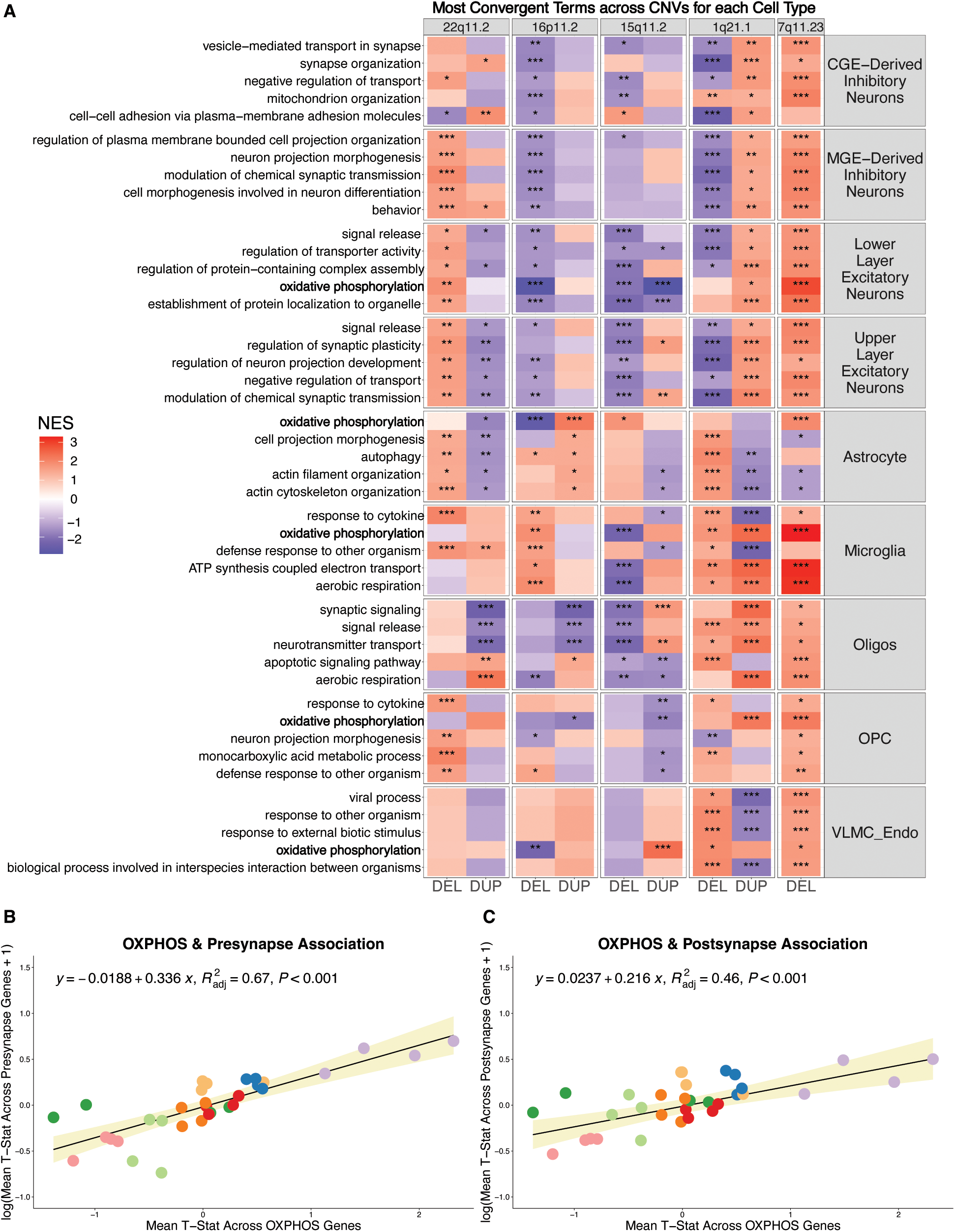
Cell type-specific enrichments across CNVs. (A) GSEA results of the top 5 most frequently implicated terms enriched at FDR <0.05 across CNVs were selected for each of the nine cell types, ordered in reverse alphabetical order. Heatmap is colored by NES, with red indicating positive NES and blue indicating negative NES. FDR significance is marked with asterisks (FDR <0.05 = *, FDR <0.01 = **, FDR < 0.001 = ***). (B) Association of OXPHOS and presynapse DGE statistics. Mean t-statistics across all genes in the OXPHOS term for each neuronal CNV model (n=36) (x-axis) and across all genes in presynapse term (y-axis), which was log-transformed for normality. (C) Association of OXPHOS and postsynaptic density DGE statistics. Same x-axis as described above and mean t-statistic across genes in postsynaptic density term for CNV models for four neuronal cell types.

We found that of all significant (FDR <0.05) pathways in the four deletion CNVs, there were 292 overlapping GOBP pathways (Figure 4A). Across all 81 models, the most frequently implicated terms included “mitochondrial ATP synthesis coupled electron transport”, “cellular respiration”, “oxidative phosphorylation” (OXPHOS), “regulation of synaptic plasticity”, and “synapse organization” (Figure 4B). Eleven of the top fifteen most frequent terms were related to two major themes: mitochondrial energy metabolism and synaptic function (Figure 4C).

Within these themes, we found that the direction of enrichment often diverged between deletions and reciprocal duplications, suggesting a CNV gene dosage effect. Among synapse-related terms, CNVs on 22q11.2 and 1q21.1 showed the most prominent directional divergence between the respective deletion and duplication: synaptic terms were positively enriched in the 22q11.2 deletion and 1q21.1 duplication and negatively enriched in the 22q11.2 duplication and 1q21.1 deletion (Figure 4D). Mitochondrial energy metabolism terms showed the most distinct reversal of direction for 16p11.2, with the deletion showing negative enrichment and the duplication showing positive enrichment of genes associated with these terms. The 7q11.23 deletion showed the strongest positive enrichment of energy metabolism-related terms across cell types of any individual CNV.

Notably, OXPHOS was the most frequently implicated term, with FDR<0.05 in 43 of the 81 models across CNVs and cell types. OXPHOS was significantly enriched in at least one cell type for all nine CNVs, and the direction of enrichment stayed mostly consistent across cell types for each CNV. The strongest OXPHOS enrichments were associated with the 7q11.23 deletion, with consistent upregulation across cell types, and the 16p11.2 deletion, with consistent downregulation across most cell types. These two deletions showed OXPHOS enrichment with an FDR<0.001 for most cell types (Figure 4E). When testing if this signal was localized to specific complexes in OXPHOS, we found that enrichment was spread across the classic electron transfer chain complexes. Specifically, the 7q11.23 and 16p11.2 deletions showed positive and negative enrichment as well as gene t-statistics for complexes I, III, IV, and V, respectively, across all neuronal and many glial cell types (Figure 4F, Figure S12).

To ensure that these findings were not a consequence of confounding or quality-related factors, we compared these results to our quality-covariate adjusted model, which accounted for RNA integrity number (RIN), mean gene detection rate, and cell count of each cell type by sample as fixed effects (Methods; Figure S7D). We observed highly correlated transcriptome-wide t-statistics between the original and the quality-covariate-adjusted model (Figure S7E). Importantly, OXPHOS enrichments remained significant and consistent across CNVs and cell types, with 38 of the 81 models meeting FDR <0.05 significance (Figure S13C), demonstrating that the OXPHOS signal was not driven by quality-related variables.

To capture the most pervasive disruptions across cell types for individual CNVs, we next examined the most consistent functional enrichments for each individual CNV (Figure S11A). Themes of synaptic disruption and OXPHOS emerged for most individual CNVs. Some CNVs, specifically the 22q11.2 duplication and 15q11.2 deletion, showed synaptic terms as the most frequently implicated terms across cell types. OXPHOS was one of the most frequently implicated terms across cell types for six of the nine CNVs. The most frequently implicated terms across cell types for the 16p11.2 and 7q11.23 deletions were related to mitochondrial energy metabolism, consistent with the strong enrichment signal for OXPHOS observed for these deletions (Figure 4E).

We also noted that a related theme of hypoxia-related terms emerged for 22q11.2 across cell types with the most frequent terms including “response to oxygen levels” and “cellular response to hypoxia” (Figure S11A). Lastly, although OXPHOS was not significantly enriched for the 22q11.2 in most cell types, another term related to energy metabolism, “glycolytic processes”, was frequently implicated across cell types (Figure S11B).

### Disruption in mitochondrial energetics extends to infant CNV carriers

We also analyzed a smaller infant dataset, which included samples from the dlPFC and ACC from infant carriers of the 7q11.23 or 15q11.2 deletions, along with matched infant non-carriers. A parallel approach for cell type annotations and clustering (Methods; Figure S6A-D), as well as DGE via dreamlet and functional enrichments via GSEA was conducted on the infant samples (Figure S6,S15). DGE models in infants showed lower residual variance, suggesting that infant brain may be less influenced by external factors than adult brain samples (Figure S14B). Genes within the deletions were downregulated, as expected. (Figure S15A). In the 7q11.23 deletion, we observed that GO terms implicated in the adult carriers were also implicated in infants, including OXPHOS (Figure S15C, D). Genes involved in OXPHOS were highly significantly (FDR <0.001) downregulated across all four neuron types in the 7q11.23 deletion in the infant brain and upregulated across several cell types in the 15q11.2 deletion (Figure S15D). Thus, while the direction of enrichment was reversed between the infant and adult samples, gene expression changes implicated OXPHOS changes in adult and infant carriers of the 7q11.23 and 15q11.2 deletions alike.

### Cell type-specific disruptions shared across multiple CNVs

Our focus next turned to identifying common signals among cell types across distinct CNVs. This analysis contextualizes alterations specific to each cell type, which is masked in bulk-tissue studies, while also revealing convergent patterns across distinct CNVs. As before, we examined the most frequent terms with FDR<0.05 within each cell type.

For neurons, we observed that the most frequently implicated terms were mainly related to regulation of the synapse, protein quality control, and cellular communication. For CGE-derived inhibitory neurons, enrichments involved cellular communication and signaling, e.g. “negative regulation of transport”, and “cell-cell adhesion via plasma membrane adhesion molecules”. MGE-derived inhibitory neurons showed enrichments in “neuron projection morphogenesis” and “modulation of chemical synaptic transmission”. Both upper- and lower-layer excitatory neurons displayed functional enrichments associated with synaptic transmission modulation, synapse organization, and mitochondrial function (Figure 5A).

For glial cell types, many of the convergent terms involved energy metabolism. The most frequently implicated enrichments for astrocytes included “actin cytoskeleton organization”, “autophagy”, and OXPHOS. Microglia-specific enrichments involved cytokine responses and terms related to OXPHOS, suggesting aberrant energy metabolism and alterations in gene expression involved in inflammation and cell survival. Oligodendrocytes and OPCs exhibited enrichments in neurotransmitter transport, cell projection morphogenesis, synapse organization, and OXPHOS, which demonstrates that the pervasive disruption of synaptic and metabolic terms across CNVs extended to major glial populations (Figure 5A). Interestingly, we observed that the directions of functional enrichments again often diverged between deletions and duplications. This was most pronounced in excitatory neurons, for example in the “regulation in synaptic plasticity” and “signal release” terms for upper-layer excitatory neurons.

Given the widespread disruptions in OXPHOS and synaptic functions across CNVs, we hypothesized that changes in gene expression for OXPHOS may coincide with changes in gene expression for the synapse in neurons. Using the SynGO database to select synapse-related terms, we found that presynapse was one of the most convergent terms across the dataset, with an FDR<0.05 in 36 of the 81 models (Figure S16A). We then observed a significant, linear relationship between the enrichment scores for OXPHOS and presynapse (Figure 5B). Postsynaptic density was also among the most convergent SynGO terms, with FDR<0.05 in 32 of the models (Figure S16A). Similarly, we observed a significant, linear relationship between the enrichment scores for OXPHOS and postsynaptic density, although this relationship was less pronounced than for the presynapse (Figure 5C).

### Mirror CNVs show gene dosage effects in synaptic and energy metabolism terms

Our collection contains four ‘mirror’ CNVs, allowing us to test gene dosage effects between deletions and reciprocal duplications. Evaluating each CNV independently, we found that several significant functional enrichments were shared but diverged in their direction for deletions and the reciprocal duplications. Previous work has demonstrated reciprocal changes in clinical phenotypes, brain morphology, and gene expression between carriers of deletions and duplications of the same genomic segments. These “mirror” phenotypes suggest gene-dosage dependent effects for these mirror CNVs across the transcriptome.

To directly test CNV gene dosage effects, we constructed gene dosage models for each mirror CNV, assigning a value of 1 for deletion carriers, 2 for non-carriers, and 3 for duplication carriers (Figure 6A). These modified DGE models test for genes that are positively or negatively correlated with CNV gene dosage for each CNV, followed by functional enrichments to evaluate terms most frequently associated with CNV gene dosage across cell types. Focusing on the larger CNVs, we show here results from the 22q11.2 and 16p11.2. The 22q11.2 CNV showed the largest number of CNV dosage-sensitive genes with FDR<0.05. (Figure 6B). In the 22q11.2 gene dosage model, like the individual CNV models, astrocytes showed the largest number of DEGs, with 607 genes at FDR <0.05, which was almost 10 times greater than the cell type with the next greatest number of DEGs (CGE-derived inhibitory neurons with 64 DEGs).Functional enrichments of DEGs detected under the gene dosage models recapitulated our previous results for individual CNVs. The 22q11.2 gene dosage model was enriched for terms related to hypoxia response and glycolytic processes, which were both negatively correlated with gene dosage. Additionally, “regulation of synaptic plasticity” was enriched in many cell types, also negatively correlated with copy number (Figure 6C). This aligns with our previous analysis of convergent synapse-related terms for the 22q11.2 deletion and duplication (Figure 4B) as well as the opposing directions shown by NES for synaptic-terms in the 22q11.2 deletion and duplication (Figure 4C).

**Figure 6:**
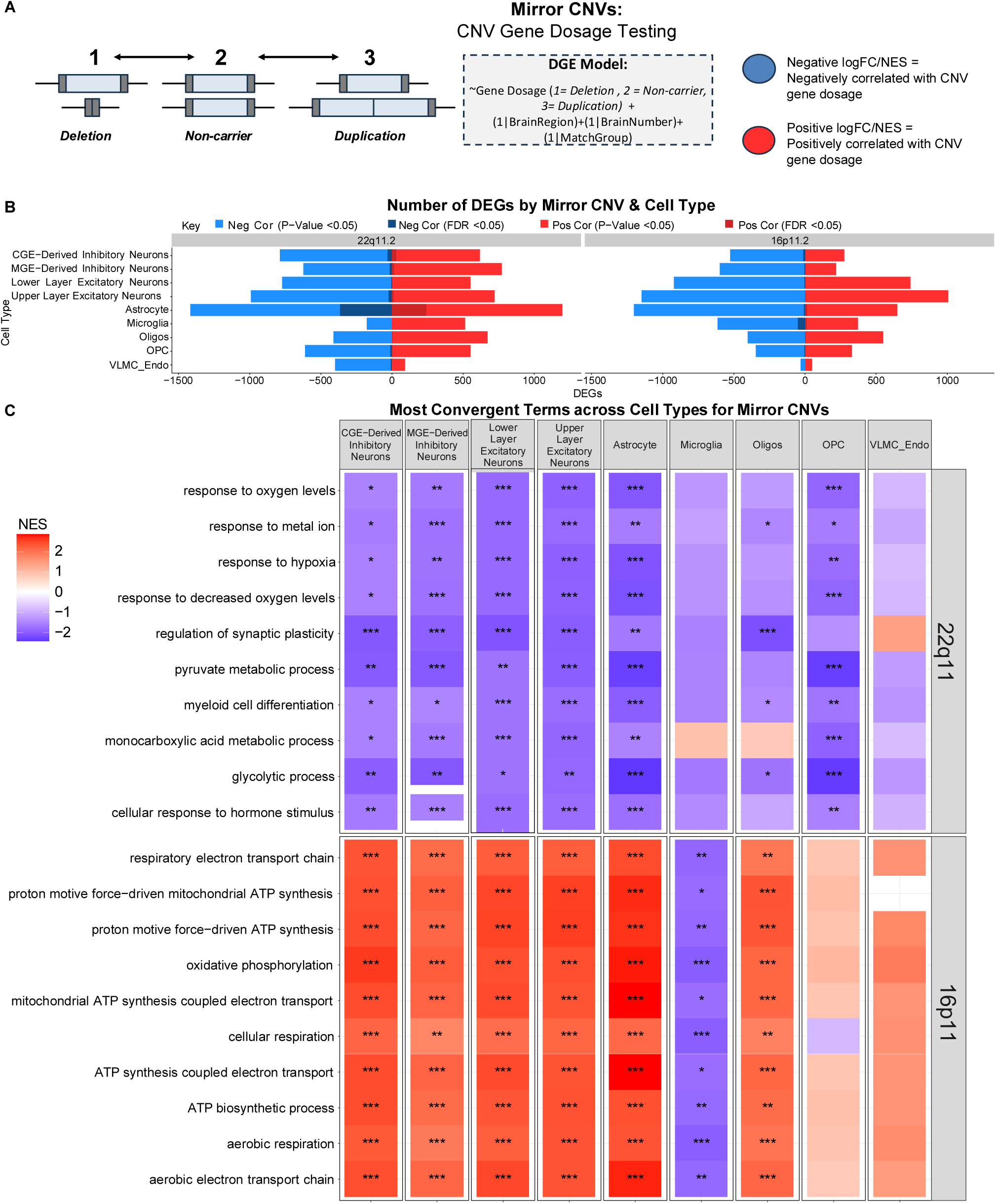
Differential gene expression testing for association with CNV gene dosage in two ‘mirror’ CNVs. (A) Schematic showing gene dosage testing. DGE was run separately for each CNV with groups of deletions, reciprocal duplication, and all non-carriers. (B) Number of DEGs (FDR <0.05) within cell types for each gene dosage model. Following the same protocol described in Figure 3A. (C) Top 10 most frequently implicated enriched terms across cell types for 22q11.2 and 16p11.2 gene dosage models are shown, ordered alphabetically.

Among carriers of 16p11.2 CNV, almost all the most frequently implicated term across cell types were related to energy metabolism: “aerobic respiration”, “OXPHOS”, “mitochondrial ATP synthesis coupled electron transport”, etc. (Figure 6C). This aligns with our previous analysis of convergent terms related to mitochondrial energy metabolism in the 16p11.2 deletion and duplication (Figure 4A). Our gene dosage model thus confirmed that OXPHOS was significantly positively correlated to 16p11.2 gene dosage for most cell types. We observed comparable levels of significance, but a negative correlation to 16p11.2 gene dosage for microglia.

## Discussion

We present a unique dataset which, for the first time, investigates the impacts of several high-risk CNVs in human brain with cell type resolution. By systematically delineating major transcriptomic effects of nine recurrent, high-penetrance neuropsychiatric CNVs across cell types, we aim to shed light on cellular pathophysiology that underlies shared phenotypes. Our study design was strengthened by matching participants based on psychiatric diagnosis and toxicology, to the extent possible, thereby controlling for treatment- and diagnosis-related epiphenomena. Leveraging single-nucleus transcriptomics, we attain cellular resolution from human brain tissue previously masked in bulk tissue studies, enabling the exploration of both individual and converging impacts of each CNV within and across cell types in the human brain. Using this resource, we discovered that differentially expressed genes in high-risk neuropsychiatric CNVs converged on mitochondrial energetics and synaptic terms. We also found cell-type specific changes, with the most pronounced effects in astrocytes and neurons. These findings provide valuable insights into the shared molecular underpinnings of high-risk CNVs, with implications for future research and therapeutic targets.

Across all DGE models, we observed a consistent upregulation for duplications and downregulation for deletions of genes within the CNV regions, affirming the reliability of the DGE models we employed. We found some disrupted gene expression in the vicinity of CNV regions, particularly in duplications, aligning with previous work finding that large CNVs can influence the expression of nearby genes (39). Despite some localized disruption, most DEGs were located outside the vicinity of the CNVs, suggesting broad transcriptomic impacts beyond their proximal effects. We found that transcriptome-wide number of DEGs detected in CNV carriers often aligned with expected levels of phenotypic risk associated with those CNVs. For instance, deletions exerted a greater influence on the transcriptome than the corresponding duplications of the same genomic region, mirroring clinical and neuroimaging studies, which have likewise found greater penetrance of neuropsychiatric disorders for pathogenic deletions than for reciprocal duplications(1,9,17,40,41). Additionally, we found that the size of the deletions CNV, as well as the summed score of the evolutionary constraint of the genes within them, was correlated with the number of DEGs. These findings were particularly evident in the 22q11.2 and 7q11.23 deletions, as they are two large deletions and showed the greatest number of DEGs. This aligns with previous work showing larger deletions to be more penetrant than smaller deletions and duplications (9), and highlights the extensive impacts of large, rare deletions on the transcriptome of human brain cell types. One exception to this was the 1q21.1 deletion, the sole carrier of which showed the highest number of DEGs despite the moderate size of the distal 1q21.1 region. 1q21.1 CNV carriers in our study varied in the involvement of the proximal TAR region, with the deletion spanning both the proximal and distal region, a larger subtype of the 1q21.1 deletion often associated with more severe phenotypes(47). Additionally, one of the three 1q21.1 distal duplications was also accompanied by a deletion in the proximal region. Due to these irregularities, the 1q21.1 CNV was not included in the CNV size and constraint analyses.

A core goal of our study was to explore the convergent effects of high-risk CNVs across cell types to provide biological insight into their shared pathophysiology. Functional enrichment analyses revealed that gene expression changes often involved functions related to energy metabolism and synaptic activity. OXPHOS was the most frequently implicated term, both with and without correction for quality-related covariates, showing the most pronounced effects in the 16p11.2 and 7q11.23 deletions. This aligns with a growing body of work implicating disruptions of mitochondrial function and OXPHOS from mouse and cellular models of the 16p11.2 deletion (48,49), 7q11.23 deletion (50,51), and 22q11.2 deletion (52–55). Importantly, some of these studies, notably Tebbenkamp et al., 2018, also completed mitochondrial assays showing aberrant mitochondrial functioning. Disruptions in mitochondrial energetics have also been observed in CNVs not covered in our study, but with similar risk profiles, for instance in cell culture models of the 3q29.2 deletion, a significant genetic risk factor for schizophrenia and bipolar disorder (56). Our findings, taken together with the aforementioned studies of individual CNVs, point to energy metabolism as a common thread among carriers of high-risk neuropsychiatric CNVs, suggesting a potential mechanism contributing to the convergent pathophysiology of these CNVs.

The second major convergent theme across all nine CNVs involved synapse-related terms, with disruptions already well-documented in several CNVs, including 22q11.2 (53,57–62), 16p11.2 (19,49,63–65), and 7q11.23 deletions (50,66) and 16p11.2 duplication (67,68). Given that synaptic activity is highly energy-demanding and relies on mitochondrial ATP production, this theme may be linked to the mitochondrial dysfunction also observed. In our data, DGE signal across neurons for genes encoding the pre-synapse and, to a lesser extent, post-synaptic density was positively correlated with those of OXPHOS, which supports this potential connection and aligns with previous works (69–74).

Using single-nucleus RNA-seq, we found that high-risk CNVs have cell-type specific effects, particularly in excitatory neurons and astrocytes. Excitatory neurons showed higher DEGs than other celltypes, with especially strong effects in the 7q11.23 deletion, consistent with prior studies highlighting its impact on this cell type (50,75). Astrocytes showed the highest number of overall and unique DEGs among all cell types, with functional convergence on stress-related terms like actin cytoskeleton organization and autophagy.

These changes suggests structural changes that may support neuroinflammation regulation, oxidative stress protection, and neuronal survival (76–79). Astrocytes also engage in essential cellular communication with neurons and are critical for supporting neuronal survival and synaptic activity (80,81). Astrocyte-specific changes in our study may reflect an adaptive response to altered neuronal energy metabolism and synaptic function, consistent with evidence of coordinated synaptic gene expression between astrocytes and neurons in schizophrenia (82). The 22q11.2 deletion exhibited markedly more DEGs in astrocytes than other cell types in both the individual CNV and gene dosage models, with significant upregulation of hypoxia-induced stress and glycolysis-related terms across astrocytes and other cells. Such changes are consistent with a shift from OXPHOS to glycolysis (anaerobic energy metabolism) in response to increased energy demand or hypoxia, a known response especially in astrocytes (83–85). This aligns with prior findings which observed altered glycolysis both in patients with 22q11.2 deletion (86) and in model systems (54), as well as with previous studies that described increased neuronal excitability (87), impaired mitochondria function (52–55,60,85), and increased glycolysis biomarkers in the 22q11.2 deletion (54,86). Together with the above-mentioned widespread changes in OXPHOS, these findings further support the notion of convergent energy metabolism changes in high-risk CNV carriers, but with a particular emphasize on glycolysis for the 22q11.2 deletion.

Despite these insights, this study has several important limitations. Despite screening more than a thousand individuals across three brain banks, the number of carriers was limited by the rarity of neuropsychiatric CNVs in the population. These results likely capture the most prominent gene expression changes; however, the sample is underpowered to reliably detect changes unique to individual CNVs. Although carriers were selected based on overlap with known recurrent high-risk CNVs, the exact breakpoints cannot be precisely determined based on SNP array data alone. While this does not affect most DGE analyses, it may complicate those that depend on gene location relative to the CNV. Although rarer cell types were present in our dataset, the DGE analysis was limited in resolution to nine major cell classes selected for frequency and even representation across the samples.

While gene expression changes in post-mortem tissue provide valuable insight, the observed changes may include both those that are caused by the CNV and those that result from the associated illnesses and treatments. By matching of carriers and non-carriers by psychiatric diagnosis and treatment, we attempted to control for those variables, but such matching is never perfect. Studies of post-mortem tissue will require further experimental validation to robustly disentangle gene expression changes caused by the CNV from secondary effects. Our study investigated transcriptomic impact of high-risk CNVs but lacked protein-level data; this would be an important next step to further understand the functional implications of these results, particularly on mitochondrial energy metabolism.

Moving forward, future research endeavors will need to functionally validate these findings of convergent mitochondrial disruptions across cell types and their connection to the synapse. Larger cohorts are essential to further validate and elucidate the complex genetic and molecular landscape of these rare, large CNVs. Additionally, it would be important to expand this work beyond the two cortical brain regions described in this study.

Neuroimaging studies have identified many changes in subcortical regions associated with CNVs (88,89), but our dataset was limited to dlPFC and ACC. Studying other brain regions could yield further insights and provide a better basis for integration with neuroimaging work, which would seek to answer how molecular changes may change brain physiology and lead to clinical phenotypes. Additionally, an avenue to further interrogate these CNVs would be through incorporating multi-omic approaches, i.e. pairing gene expression with other genomic assays. This future work will provide a deeper understanding of CNV biology and the relationship between mitochondrial and synaptic dysfunction, paving the way for the development of biomarkers, and more effective diagnostic and therapeutic strategies for individuals with high-risk neuropsychiatric CNVs.

## Acknowledgements

We would like to thank the families who donated the brains of their loved ones for research, Maree Webster, Tom Blanchard, Robert Johnson, and the NeuroBioBank team for brain banking resources, dissection, and distribution of brain tissue from the Stanley Medical Research Institute and the University of Maryland Brain and Tissue Bank, Barbara Lipska, Ajeet Mandal, Bhaskar Kolachana, and Vesna Imamovic for brain banking resources and assistance with genotyping and brain tissue dissection at the NIMH Human Brain Collection Core, Yunlong He, Bao Tran, and other members of the CCR Sequencing Facility for nuclei isolation, library preparation, and sequencing, Asya Khlebodorova and Justin Lack for assistance with bulk RNA-seq data processing, all members of the Human Genetics Branch and Human Brain Collection Core (HBCC) for frequent discussions and ideas, and the NIH Biowulf team for providing computational resources. This project is supported by the NIMH Intramural Research Program: ZICMH002903 (Human Brain Collection Core) and ZIAMH002810 (Human Genetics Branch).

## Author contributions

GD analyzed data and wrote the paper. NA performed CNV calling. NA, SM, PKA, FJM, and AS participated in study design and coordinated data acquisition. PKA performed and coordinated dissections. QX extracted RNA and measured RIN. QX, NF, and BK performed genotyping. NA, SM, PKA, AR, SL, AD, XJ, MDG, KFB, GH, PR, FJM, and AS assisted with data analysis and interpretation. FJM and AS assisted with writing the paper. All authors provided feedback.

