## Supplementary Methods for "Transcriptomic signatures of nine high-risk neuropsychiatric copy number variants in human brain cells"

*Denotes equal contribution

This PDF Includes:

- Supplementary methods
- Figure S1-16
- Table S1
- Legends for Table S2-9

### Supplementary Methods:

CNVs of interest:

We only included CNVs with high-confidence CNV calls, in loci with high penetrance for neuropsychiatric conditions [1,2], and with at least two individuals harboring a CNV (deletion or duplication) in the same locus. We identified 15 CNV carriers who met these conditions. They included reciprocal deletions and duplications on chromosomes 22q11.2 (VCFS, proximal, A-D), 16p11.2 (proximal, BP4-BP5), 15q11.2 (BP1-BP2), 1q21.1 (all covering BP3-BP4, with variable involvement of the proximal BP2-BP3 TAR locus), and 7q11.23 (WBS) deletions.

Matching approach and demographics

Each individual harboring a known high-risk neuropsychiatric CNV, as defined above, was matched as close as possible to at least two presumptive non-carrier individuals from the same brain bank with similar demographic characteristics, based on sex, ethnicity, age of death, psychiatric diagnosis, and drug profile in toxicological screening (when available). In cases where two CNV carriers had very similar demographic features, the same non-carrier individuals served as matches for both. This matching approach created groups of three or more individuals with similar characteristics, which we refer to as 'match groups'. Demographic variables across the dataset are summarized in Table S3. Since we are not permitted to publish individual-level metadata, only summary-level information is provided. The psychiatric diagnoses among the CNV carriers included non-psychiatric controls (n=3), major depressive disorder (n=3), bipolar disorder (n=2), schizophrenia (n=1), autism spectrum disorder (n=1), eating disorder (n=1), Williams Syndrome (n=2) which reflects the composition of the screened population, prevalence of these CNVs in the population, and their pleiotropic nature[2–4]. Serious mental illness and its treatment are known to have an impact on the transcriptome[5]. To increase the chances of the transcriptomic effects being driven by the CNV rather than the psychiatric condition or associated epiphenomena, our matching approach included psychiatric diagnosis and toxicology, whenever possible, in addition to standard demographic matching on age, sex and ethnicity.

Regions of interest

Whenever available, dorsolateral prefrontal cortex (dlPFC) and anterior cingulate cortex (ACC) were dissected. 50 mg of frozen tissue were used for nuclei isolation and single-nucleus RNA-sequencing, while another 50 mg were used for RNA isolation, followed by bulk RNA-sequencing and measurement of RNA integrity numbers (RIN).

#### Quality Control, Nuclei Filtering, and Clustering

All analyses were performed with R version 4.2.2. Data from adult (>10 years old) and infant (<1 year old) brain samples were processed separately. Starting with the adult data, we converted the single-nucleus RNA-sequencing counts into a Seurat object, which we then iteratively filtered and visualized with uniform manifold approximation and projections (UMAPs). To ensure data quality and reliability, we implemented a series of rigorous quality control and filtering steps (Figure S2). Utilizing SoupX (v 1.6.2) [6], we eliminated ambient RNA using the default values provided by the software.

Sample outliers were determined by Pearson correlation analysis of the normalized count matrix after pseudobulking across the 36,601 mapped genes. We removed two non-carrier samples (26 and 67) as they show correlation values ≤0.7, the lowest overall correlations compared to the other 67 samples (Figure S3A).

We employed scDblFinder (v 1.12.0) [7] to identify and filter out doublets in our data. scDblFinder was run on each sample using the recommended default values. We did not observe a noticeable concentration of doublets in any cluster (all <50% doublets) (Figure S2A). We thus removed any cell classified as a doublet according to scDblFinder.

We established the following criteria for initial nuclei inclusion: nuclei with >200 unique genes detected, > 500 number of transcripts, and a fraction of mitochondria-encoded genes (percent.mt) <5.0% (Figure S2C). Using the Seurat package (v 4.3.0) [8], the data underwent normalization, dimensionality reductions, and unsupervised clustering with Louvain clustering resolution of 1.0.

#### Harmony Integration and Additional Nuclei Filtering

To address potential batch effects biasing clustering across the dataset, we applied the Harmony algorithm (v 0.1.1) [9]. We used Harmony to control for match group, as it discerned cell type clusters with the best resolution. This was expected, since match groups were designed to control for the commonly observed confounding effects of sex, ethnicity, psychiatric diagnosis, and age. Utilizing match group as our integration variable with 50 dimensions, we improved cluster resolution between distinct cell types.

However, we still encountered noticeable bridging between major cell classes in the UMAP visualization, suggestive of persistent debris or poor-quality nuclei. The bridged clusters were also the clusters with the highest percent.mt suggesting that this bridging was an artifact of debris with high mitochondria content. We conducted quality filtering and Harmony integration by removing clusters with highest mean percent.mt. We started with clusters exceeding 2.5%, then removed clusters with percent.mt exceeding 2.0% and reran Harmony after both steps. Rigorous iterations of cluster removal by percent.mt filtering allowed for fewer bridged clusters and better cell type resolution (Figure S2B).

#### Cell Type Annotations and Sample Filtering

We utilized multiple approaches to annotate cell types and checked for consistency across approaches. First, we explored the expression of top marker genes associated with expected cell types. Then we employed Azimuth software (v 0.4.6) to annotate cell types, using the human motor cortex data set from the Allen Institute of Brain Research as a reference [8,10].

In picking cell type resolution, we aimed to achieve a high level of biologically meaningful resolution, while ensuring robust representation across all samples. Cells were annotated by identifying the most predominant cell type with each cluster from Seurat, based on subclass provided by Azimuth. Although Azimuth allowed us to resolve cell types in fine detail, we choose to limit our analysis to nine broad cell types based on cell type counts across samples, grouping some of the smaller populations of neurons and glia. (Figure S4A). We settled on nine well-represented cell types: astrocytes, microglia, oligodendrocytes, OPCs, vascular leptomeningeal cell (VLMC) and endothelial cells, upper- and lower-layer excitatory neurons, and medial ganglionic eminences (MGE)- and caudal ganglionic eminences (CGE)-derived inhibitory neurons (Figure 1E). VLMCs and endothelial cells were grouped together as they both had very low cell counts and serve similar biological functions. Low cell counts of VLMC and endothelial cell types are not unexpected in dlPFC and ACC[11]. We compared these annotations to marker gene expression to ensure confidence (Figure S4 A-C). We observed similar proportions of the nine broad cell types between dlPFC and ACC, after normalizing by the number of samples from both brain region (Figure S5A,B), consistent with prior work [12].

After classifying cell types, we filtered out samples with low neuron numbers. We removed sample 30, which contained fewer than 20 neurons across all four neuron types (Figure S3C; 5C). Following the same criteria for outlier detection as used before, we analyzed Pearson correlations between samples on the normalized pseudobulked expression matrix across the 36601 genes (Figure S3B). Sample 30 also had low correlations across the 67 samples and, along with the underrepresentation of neuronal cell types, was thus excluded on this basis. After removing the outlier sample, each cell type had greater than 10 cells in each sample, except VLMCs.

#### Testing of Cell type Proportion

To rule out major cell type shifts and detect potential biases in the relative cell type abundance, we calculated the proportion of each cell type using the total cell count for each sample as the denominator and respective cell counts for each of the nine cell types as the numerator. Within each cell type, we ran a linear mixed model using lme4 (linear mixed-effects models) [13] procedure following the formulas:

*(1) celltype_proportion~CNV +(1|BrainRegion)  (1 | BrainNumber) +(1|Match_group)*

*(2) celltype_count~CNV +(1|BrainRegion) +(1 |BrainNumber) +(1 |Match_group)*

We used square root and log transformations on the cell type proportion and counts respectively for the model to meet normality. Transformation for each cell type and model were chosen based on the Shapiro-Wilks test, requiring a p-value >0.05 of the residuals. We tested for the effect of CNV with match group, brain number, and region as random effects, since they are nested in the study design. We did not observe significant cell type shifts across most CNVs. One exception was the 15q11.2 deletion, which showed significant differences in cell type proportion for most cell types. The cell type counts for the 15q11.2 deletion, however, only showed significantly higher numbers of microglia and oligodendrocytes. These two cell types likely drive the significant results across proportions. Additionally, OPCs in the 22q11.2 deletion were significant for both cell type proportion and counts, with fewer OPCs in carriers than non-carriers (Table S 5;6). Given the limited sample size, we cannot conclude if differences in cell type proportion were due to biological or technical factors. However, the involvement of oligendrocytes, OPCs, and microglia suggests that they may be attributable to the ratio of grey and white matter in the dissection.

Differential Gene Expression:

We ran DGE using dreamlet (v 0.99.6), because it has the capability to use mixed models to addresses the nested nature of our data. Dreamlet also adopts a pseudobulking strategy for single-nucleus data, which has shown improved performance compared to single-cell approaches[14]. Dreamlet incorporates two-levels of precision weights, accommodating variations in both cell count and sequencing depth. This was beneficial to our study design to account for the observed variability in cell type counts described above.

We followed default settings in dreamlet for gene and sample filtering, which requires a minimum number of five reads for a gene to be considered expressed in a sample, minimum of four samples passing cutoff for a cell type to be retained, and minimum of 40% of retained samples to have non-zero counts for a gene to be retained.

Utilizing matched non-carriers enabled us to account for known confounders despite limited sample size and better target transcriptomic changes driven by each CNV. However, it was important to delineate how well our non-carrier controls were matched and determine correction for additional covariates as needed. We ran variance partitioning and linear regressions to decide what other covariates to include and found none retained significance across all nine celltypes. However, for comparison, we conducted a parallel analysis with additional quality-related covariates.

The adjusted model was:

*~CNV + (1|BrainRegion) + (1|BrainNumber) + (1|MatchGroup) + RIN + nFeature + Cell.Type.Counts*

We observed high correlations of t-statistics between the original model and the quality-covariate adjusted model, except for oligodendrocytes in the 15q11.2 deletion. This was consistent with the significant differences in cell-type proportions in this sample, which had suggested more white matter in that dissection (Figure S7E). Given that this is likely driven by dissection, we believe the 15q11.2 oligodendrocyte results should be interpreted with caution.

Model Selection:

Using variance partitioning, we assessed correlations among the covariates and found that match group was significantly correlated with age, sex, ethnicity, etc. as expected (Figure S7A). Exploring the variance explained by each covariate indicated that CNV status and match group account for the largest amount of variance (Figure S7B).

We also performed principal component analysis and used linear regression to test for association of known covariates with the top principal components (PC), finding that match group was the most significant covariate (Figure S7C).

After residualizing for match group, brain region, and subject ID as random effects, we found no covariates were significantly associated with PC1- at p-value < 0.05 in all cell types (Figure S7D).

Some quality-related variables were significant in several cell types, such as cell type count. While dreamlet corrects for differences in cell type counts with precision weights, we created a more rigorous quality-covariate adjusted model for comparison to ensure differences in cell type count and sample quality between carriers and non-carriers did not influence DGE results. We selected quality covariates with a p-value < 0.01 in PC1 for at least one cell type to be added to a quality-covariate-adjusted model. Three covariates met this criterion: RNA integrity number (RIN), mean number of genes per cell by sample (nFeature), and cell count of each cell type by sample.

Infant data:

The infant data underwent the same pipeline as the adult data. Moreover, the same QC steps were applied to the infant samples as with the adult samples, with Harmony integrations and removal of clusters with the highest percent mt. The nuclei were then annotated for cell type using Azimuth software (v 0.4.6) [8,10], with the human motor cortex data set from the Allen Institute of Brain Research as a reference [8] (Figure S6A-D). The only change in cell type classification was that oligodendrocytes and OPCs were merged into a single category labeled OPC_Oligo, due to the low number of oligodendrocytes in infant brain samples and the biological similarities between the two cell types.

We followed the same analysis pipeline as for the adult dataset for differential gene expression with exploration of metadata by variance partitioning and linear regression of covariates (Figure S14 A, B). The final model stayed consistent with the adult dataset as:

$$\sim CNV + (1|BrainRegion) + (1|BrainNumber) + (1|MatchGroup)$$

Similarly, the default filtering values were retained from dreamlet for the infant dataset. Functional enrichments were achieved using fGSEA [15] on the GO BP and MitoCarta database[16].

LOEUF Scores

To characterize genes within the CNV regions, the loss-of-function observed/expected upper fraction (LOEUF score) was used as a metric for risk. LOEUF scores are a continuous metric to quantify the intolerance of a gene to a loss of function variant and were obtained from the GnomAD database [17] (v4.1.0). To aggregate constraint scores for each CNV, inverse LOEUF scores (1/LOEUF) were summed for all CNV genes with available constraint metrics for each DGE model, as previously described [3]. A higher inverse-LOEUF score indicates intolerance to loss of function and suggests that the gene is essential for normal function and survival.

Functional enrichments:

Rank-based functional enrichment testing approaches were used to mitigate noise and unreliable results due to arbitrary p-value thresholds, known to influence threshold-based functional enrichment testing results [18,19]. GSEA was our primary functional enrichment testing method as it is widely used and acknowledged to be a strong candidate for discovering biological insights for RNA-seq data analysis [18,20].

We tested for functional enrichments using Gene Ontology (GO) Biological Processes (BP) database [21–24]. Mitocarta [16] was used to obtain more detailed curated terms related to mitochondrial functions, including OXPHOS complexes. We also used SynGO to select terms related to the synapse [25].

We applied a filtering criterion for functional enrichments, retaining only those with false discover rate (FDR) < 0.05 as a threshold for significance. To assess convergence across the dataset, we explored terms with the highest overlap in significant terms across both CNVs and cell types, in all 81 models (DGE across

9 CNVs and 9 cell types). Any term with the same intersection size was then ranked by FDR for selection of top terms. To explore direction of enrichment, we utilized the normalized enrichment score (NES). This value accounts for the differences in gene set size.

Additionally, we explored convergence at the level of the CNV and at the level of the cell type, by extracting terms with the highest intersection size (1) across cell types for each CNV and conversely (2) across CNVs within each cell type. When terms had the same number of intersections, they were ranked by decreasing summed FDR value.

### Supplementary Figures:

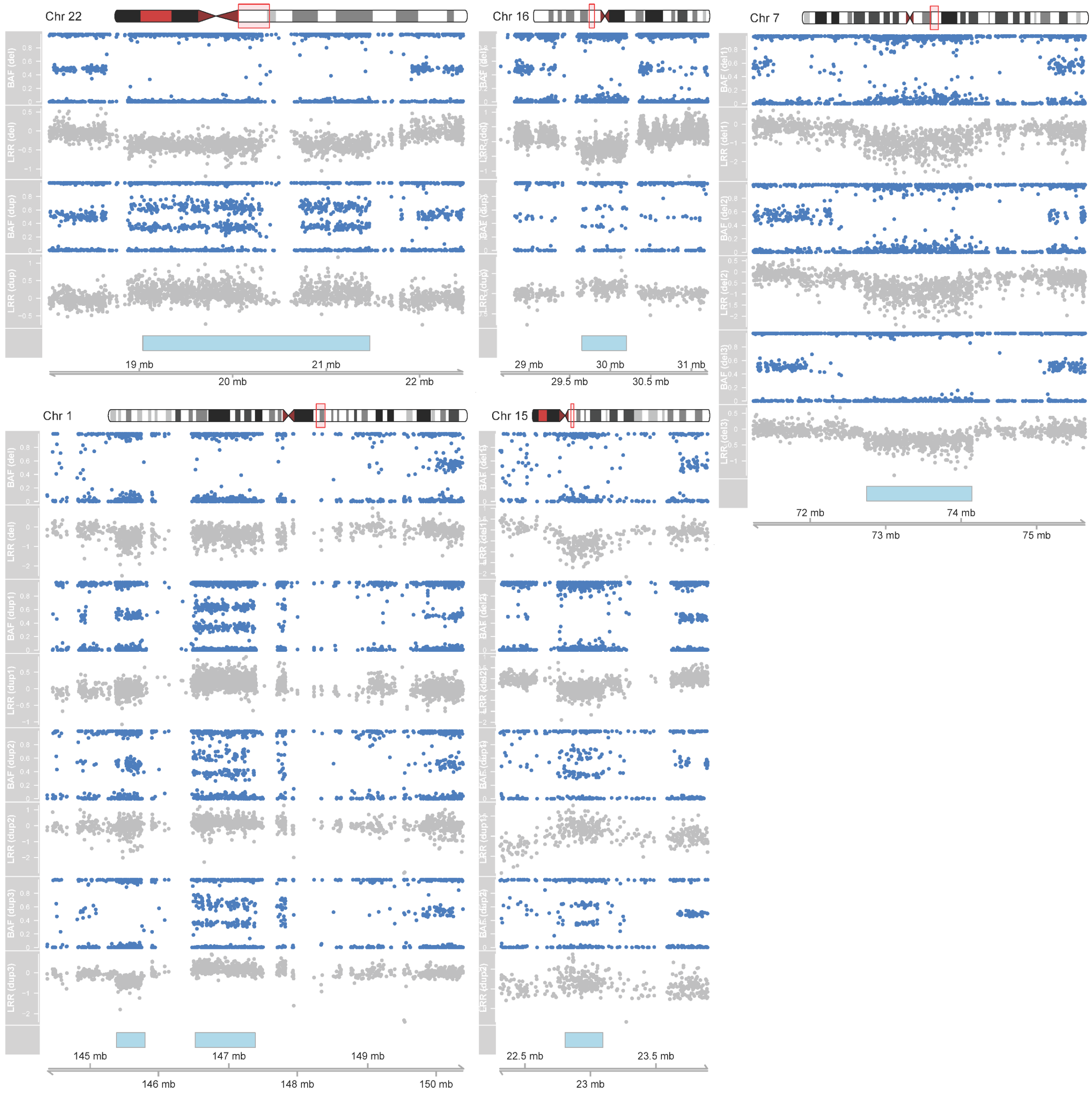


***Figure S1: Ascertainment of copy number variants.*** *B allele frequencies (BAF, blue dots) and Log R ratios (LRR, grey dots) from the raw Illumina arrays are shown for each of the 15 high-risk CNV carriers: (A) One deletion and one duplication in the 22q11.2 region (associated with DiGeorge/Velocardiofacial Syndrome [VCFS], proximal, A-D). (B) One deletion and one duplication in the 16p11.2 region (proximal, BP4-BP5). (C) Three deletions in the 7q11.23 region (associated with Williams-Beuren Syndrome [WBS]) including one infant case (del1). (D) One deletion and three duplications in the distal 1q21.1 region (BP3-BP4) with variable involvement of the proximal locus (BP2-BP3, associated with Thrombocytopenia and Absent Radis [TAR] Syndrome), with the deletion spanning both loci and one of the three duplications (dup3) in the distal locus with a deletion of the proximal locus. (E) Two deletions and two duplications in*

*the 15q11.2 region (BP1-BP2), including one infant (del1). Chromosomal locations are based on GRCh37 coordinates with reference CNV coordinates (light blue bars) derived from ClinGen[1].*

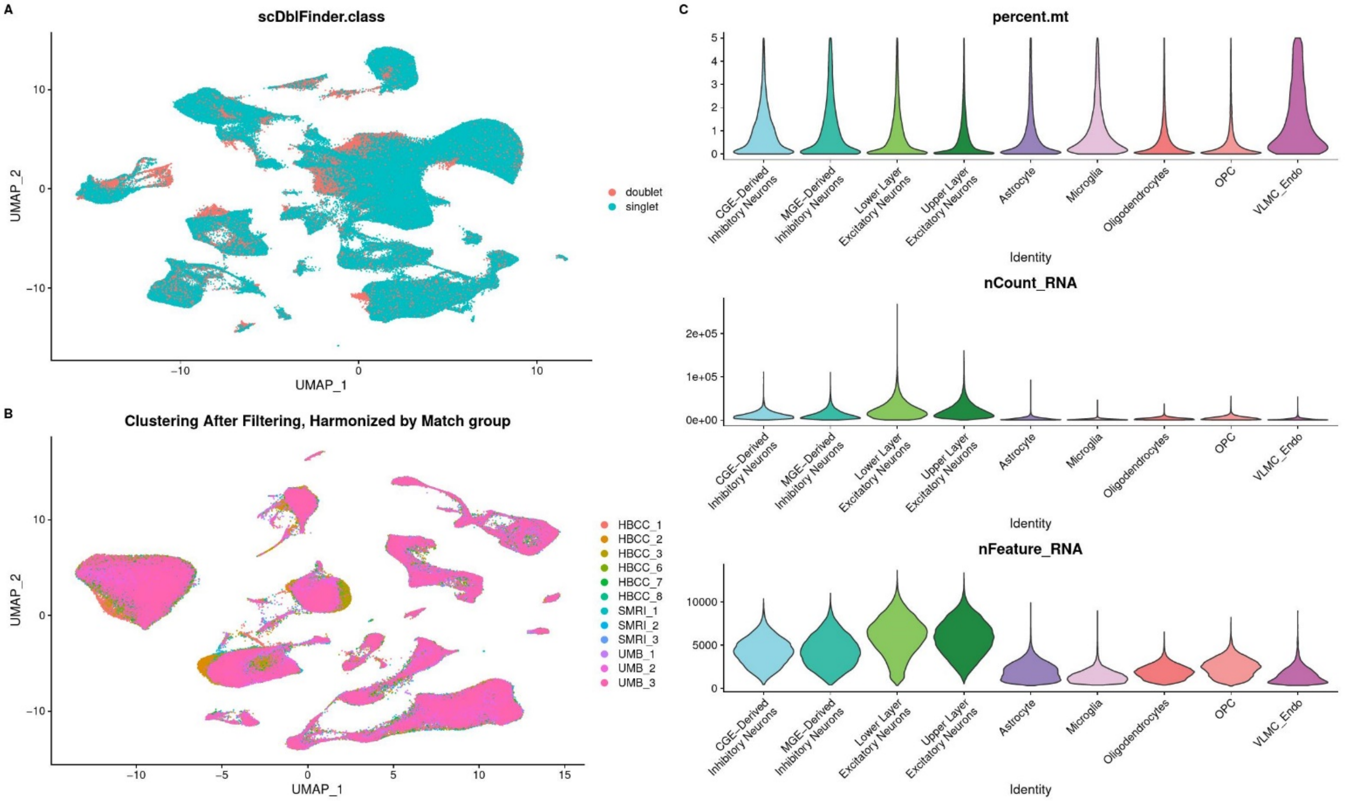


***Figure S2: Unsupervised clustering and quality control (QC****). (A) Uniform manifold approximation and projection (UMAP) visualization of interim Seurat[26] object, illustrating classification of doublets and singlets using scDblFinder[7]. Doublets were removed from downstream analysis. (B) UMAP visualizations of filtered Seurat object colored by match group (which involves sex, ethnicity, age, and psychiatric diagnosis) after Harmony integration. (C) Violin plot of mitochondria-encoded transcripts fraction per cell (percent.mt), number of transcripts within a cell (nCount), and number of unique genes detected per cell (nFeature), grouped by cell type.*

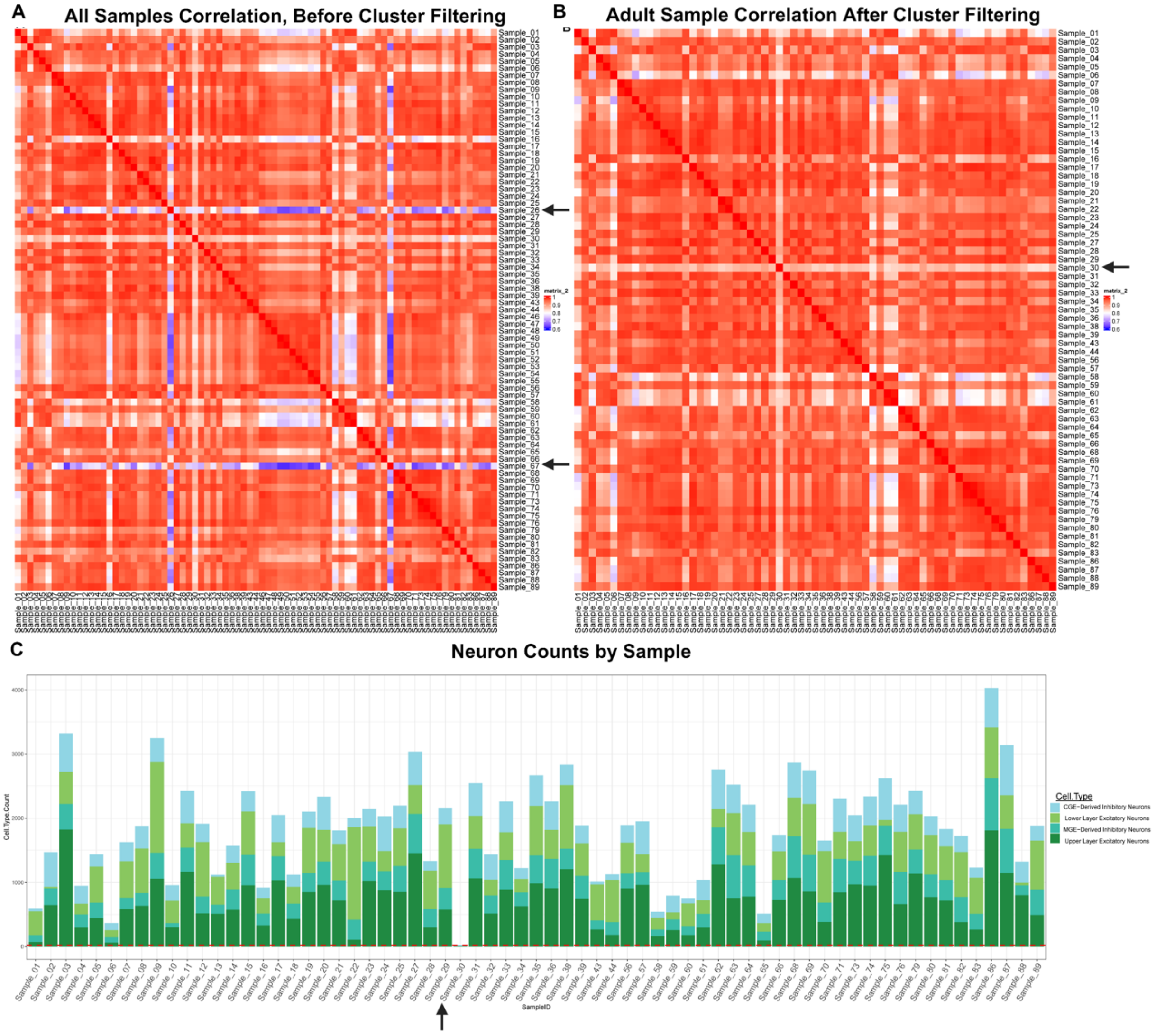


***Figure S3***: ***Sample outlier detection and filtering***. *(A) Pearson correlations between samples of pseudobulked normalized count matrix across 36601 genes before separating adult and infant samples or any downstream filtering of clusters. Sample 26 and 67 showed the lowest correlations (>0.70) and were removed from downstream analyses. (B) Pearson correlations of pseudobulked normalized count matrix of adult brain samples across 36601 genes after nuclei QC and filtering. Sample 30 shows low correlations across samples, including those within the same match group. (C) Neuron counts across adult samples. Sample 30 showed fewer than 20 total neurons and as shown in B showed consistently low Pearson correlations across the other 66 samples, with the second lowest summed correlation across all samples. Sample 30 was therefore removed from differential gene expression analysis.*

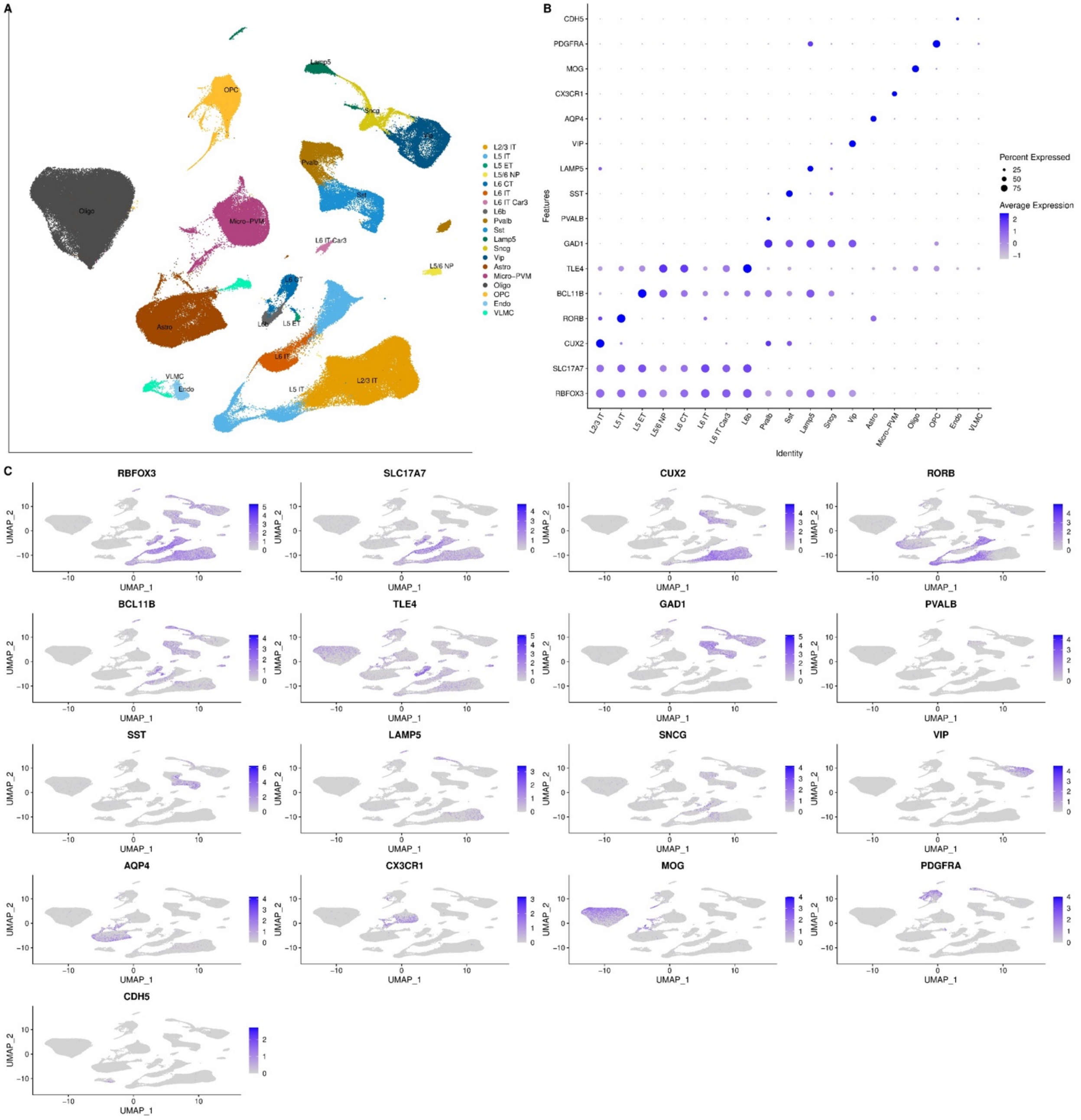


***Figure S4*: *Cell type annotations.*** *(A) UMAP visualization colored by annotated clusters using the "subclass" annotation determined by Azimuth[8] based on the Allen Institute Motor Cortex[10] reference dataset. The subclass that described the greatest number of cells in each cluster was chosen. Broader neuronal cell type annotations were defined by grouping PVALB and SST inhibitory neurons as medial ganglionic eminence (MGE)-Derived Inhibitory Neurons, while LAMP5, VIP, and SNCG inhibitory neurons were grouped and annotated as caudal ganglionic eminence (CGE)-Derived Inhibitory Neurons. Excitatory neurons from layers 2 and 3 were annotated as Upper Layer Excitatory Neurons and those from layers 5 and 6 were annotated as Lower Layer Excitatory Neurons. (B) Normalized expression dot plot of marker genes across broad cell type annotations. (C) UMAP visualization colored by expression of marker genes for major neuronal and glial cell types.*

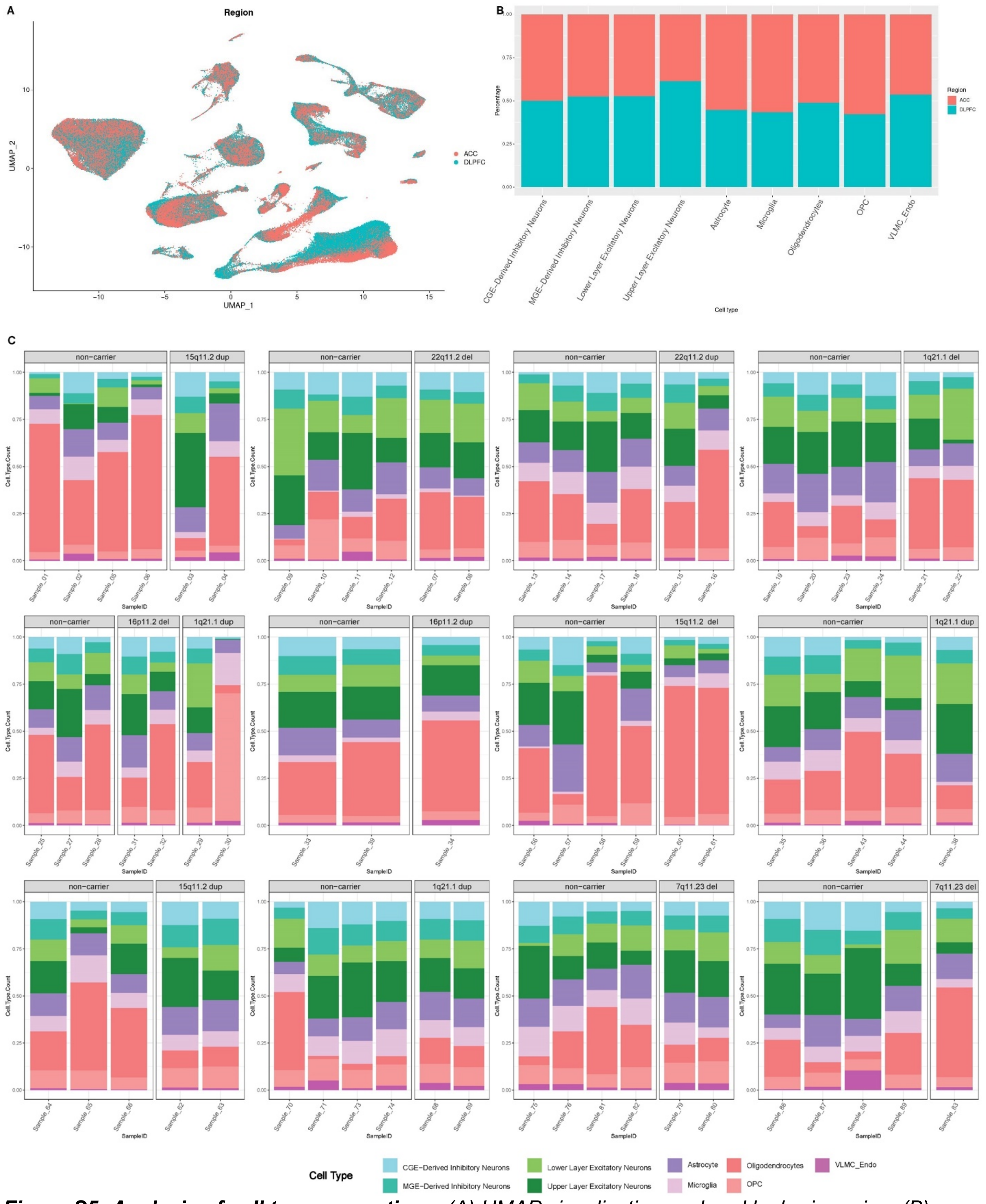


***Figure S5: Analysis of cell type proportions.*** *(A) UMAP visualizations colored by brain region. (B) Mean nuclei count of each cell type by region, after normalizing by number of samples for each region, showing roughly even distribution of major cell types across dlPFC and ACC. (C) Cell type proportions across the 67 adult brain samples, split by match group to compare carriers with matched non-carriers. Sample 30 was removed due to low neuronal counts (see Figure S3). Brain samples from the 15q11.2*

*deletion carrier showed higher proportions of oligodendrocytes compared to non-carriers, possibly due to dissection variability.*

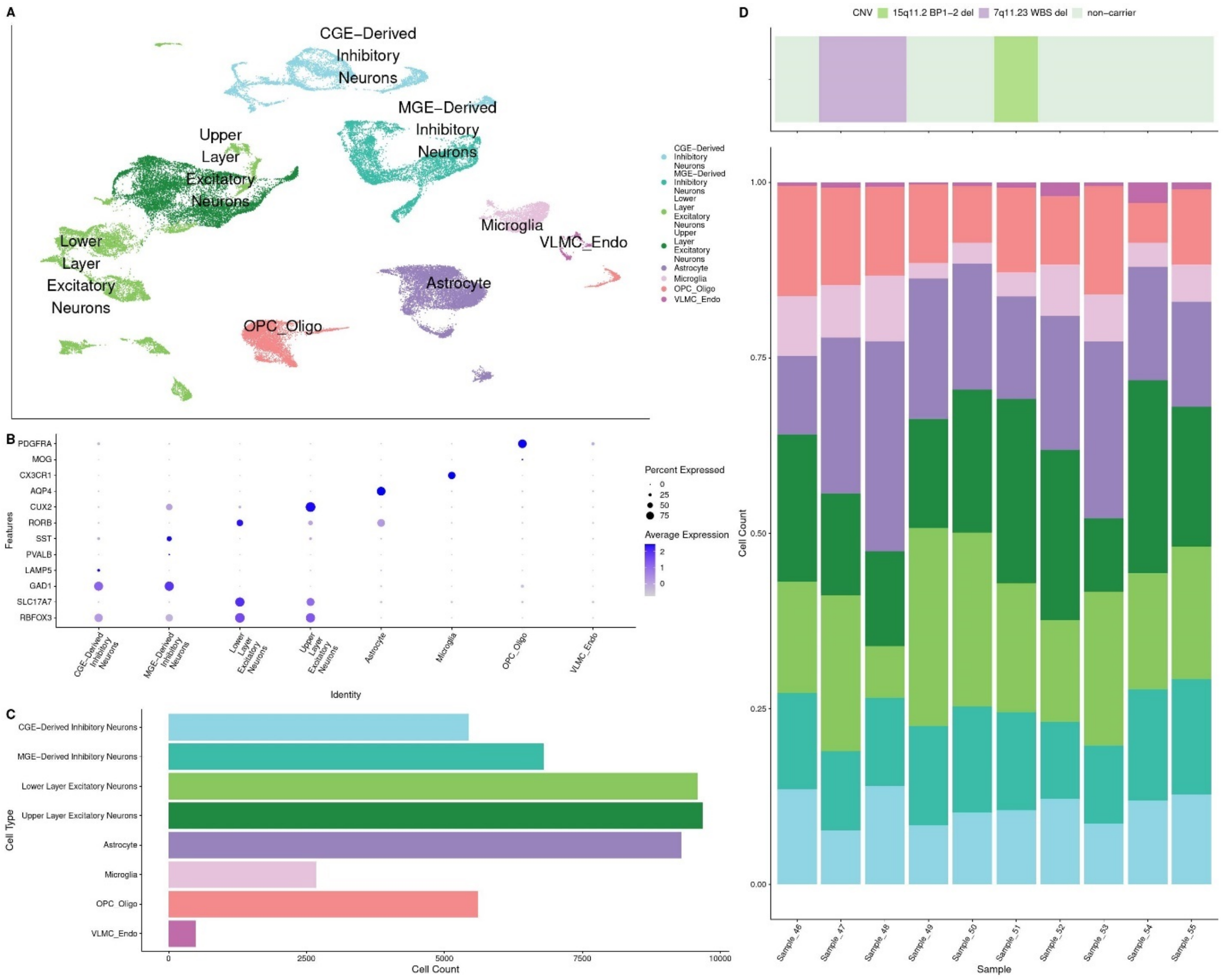


***Figure S6: Infant data clustering and QC**. (A) UMAP visualization of 49581 cells colored by cell type, annotated in the same manner as in the adult samples (see also Figure S3). OPCs and oligodendrocytes are grouped together due to low oligodendrocyte counts in infant brain samples. (B) Dot plot of normalized expression for top marker genes of the nine annotated cell types. (C) Barplot depicting nuclei counts by cell type across the ten infant brain samples. (D) Stacked barplot depicting cell type proportion by sample with color bar annotating CNV.*

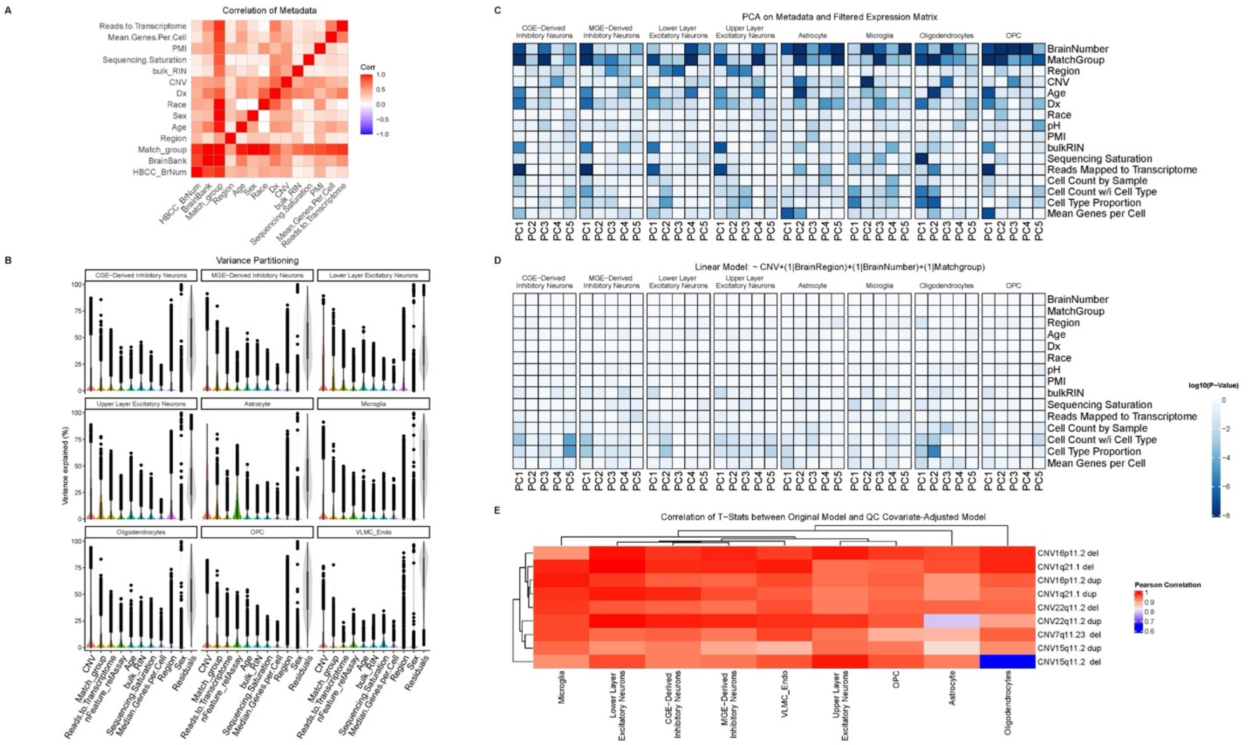


***Figure S7**: **Covariate selection for adult CNV dataset.*** *(A) Canonical Correlation Analysis of covariates using canCorPairs function of the variance partitioning[27] package. (B) Variance components analysis from variance partitioning package for known covariates shows CNV carrier status and match group as the major sources of variance across cell types. Sex accounts for a small amount of total variance, but a few genes, such as XIST and Y-chromosome genes, are strongly associated with sex, as expected. (C) Heatmap showing relationship of top five principal components (PCs) and known covariates. PCs are derived from principal component analysis (PCA) of the pseudobulked transcriptome for each cell type. Colors on the heatmap represent the log-transformed ANOVA F-test p-values. (D) Heatmap showing the relationship between residualized PCs and known covariates, where the same top five PCs as in C are corrected for CNV +(1|BrainRegion) +(1|BrainNumber) +(1|MatchGroup) and tested against each known covariate, with colors representing log-transformed ANOVA F-test p-values. (E) Heatmap showing Pearson correlation of transcriptome-wide t-statistics from differential gene expression between original model and QC-covariate adjusted model, accounting for three additional covariates, (See also Methods section* Differential Gene Expression*) across nine cell types (x-axis) for each of the nine CNVs (y-axis).*

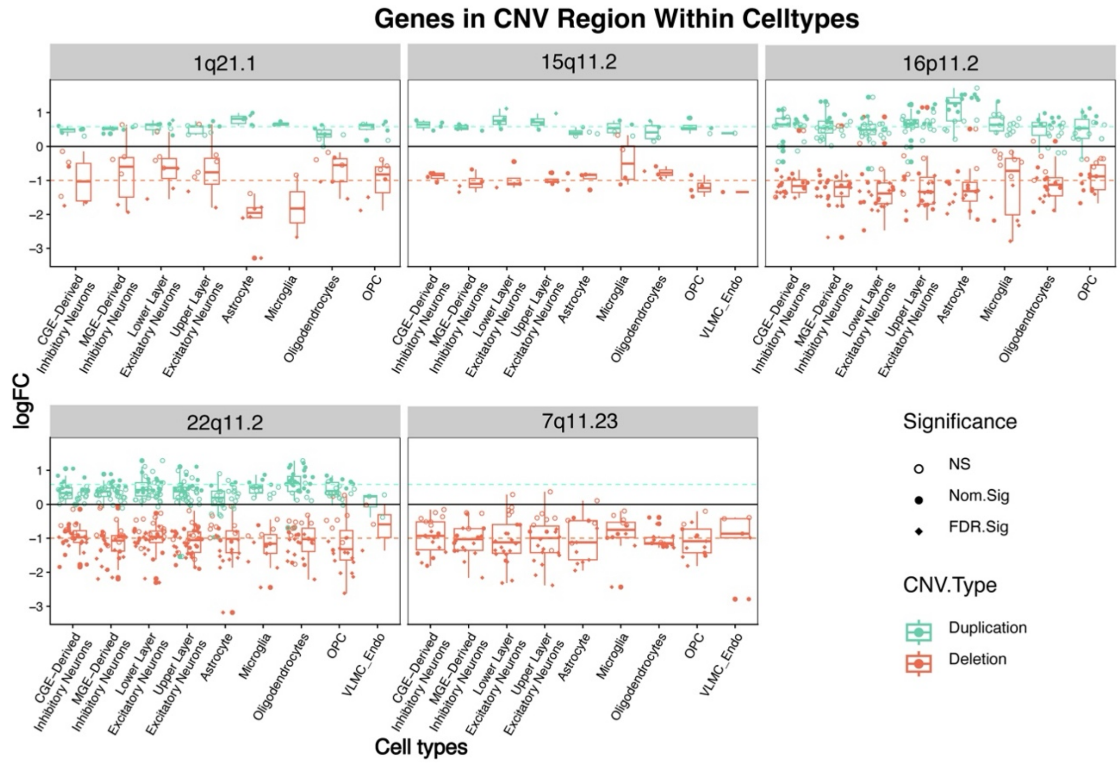


***Figure S8**: Differential gene expression results within CNV regions for each cell type.* *Boxplot with points showing logFC of individual CNV genes (see also Table S1, S2) for each cell type. CNV genes were obtained using same coordinates described in Figure 2. Dotted lines denote expected logFC for deletions (-1) and duplications (0.585) based on gene copy number.*

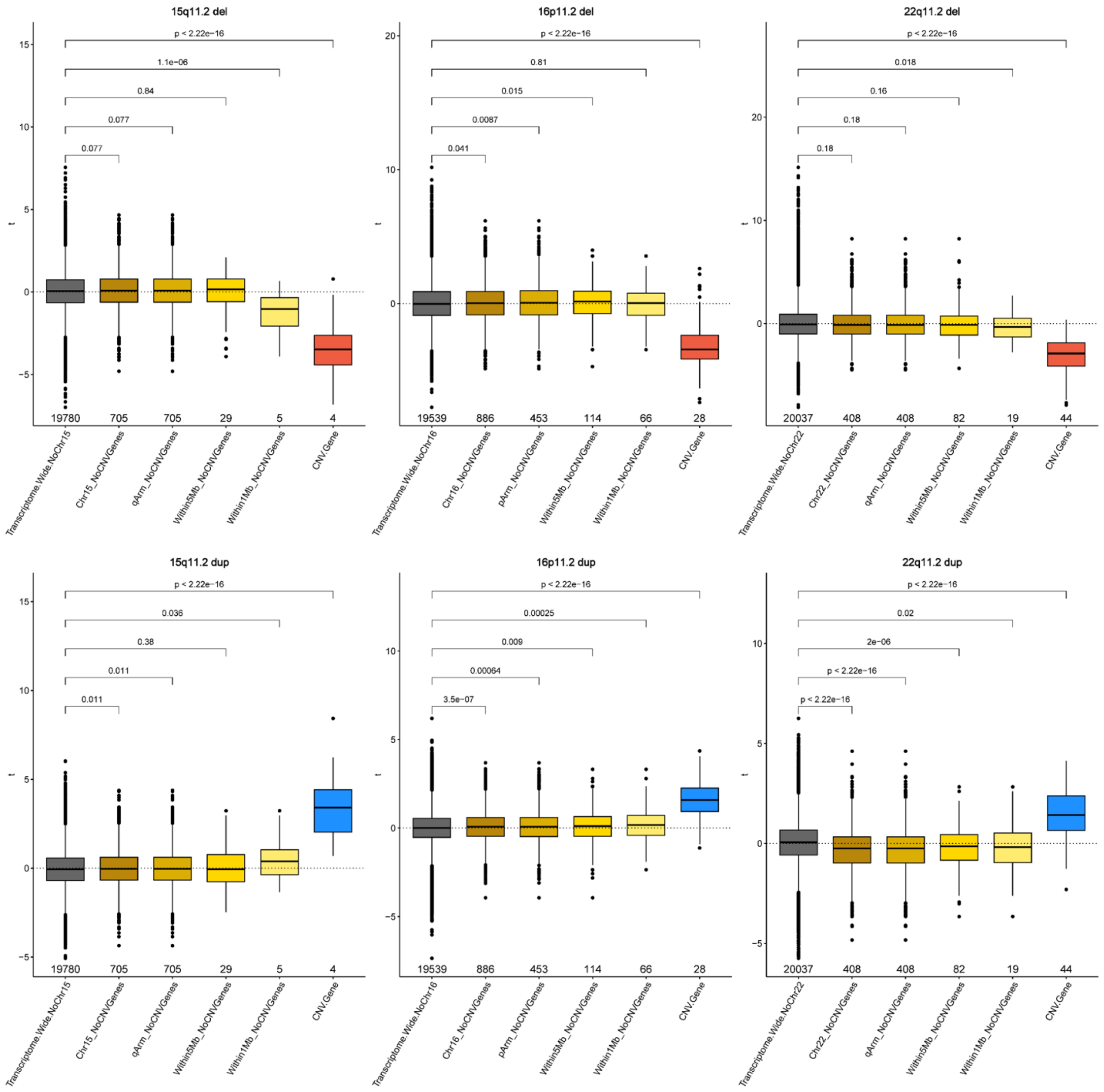


***Figure S9: Differential gene expression of genes based on genomic location respective to CNV region for mirror CNVs.*** *Boxplot showing t-statistic (y-axis) from DGE for genes within each CNV region and genes outside of each CNV region after excluding the genes within the CNV: within 1 Mb of the CNV, within 5 Mb of the CNV, within the arm of the CNV chromosome, on the CNV chromosome, and transcriptome-wide. Each dot represents a gene from the CNV DGE model for each cell type, so genes are repeated for each cell type they met filtering criterion for. P-values are based on two-sided Mann-Whitney U-test of differences of t-statistic for genes in proximity to CNV region and across the transcriptome. The number of unique genes in each category is listed above the x-axis labels.*

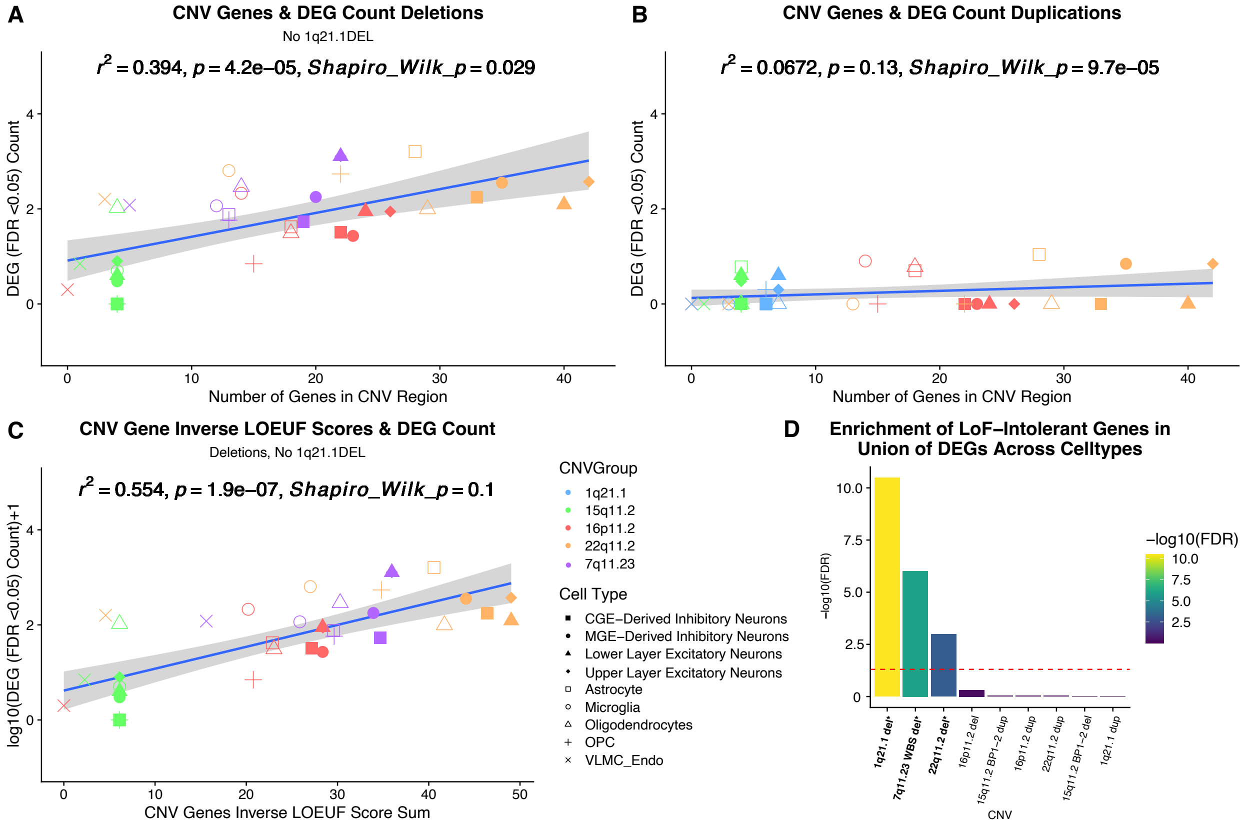


***Figure S10: Relationship between CNV size and the extent of transcriptomic disruption.*** *(A) The number of DEGs (genes that meet FDR <0.05; y-axis) shown as a function of the number of CNV genes (after gene filtering by expression levels; x-axis) for deletion CNVs. (B) Same plot as in A but for duplications. The y-axis was log-transformed to improve the Shapiro-Wilk testing of residuals normality assumptions. Deletions show a significant positive association, whereas duplications show no association. (C) Association between the inverse LOEUF score sum of CNV genes and the DEG count for deletions, which show significantly positive association. CNVs in the 1q21.1 region were excluded from all analyses of association with CNV genes, due to variable involvement of the proximal TAR region, as well as the deviations from expectations with the large number of DEGs for the 1q21.1 deletion. (D) Chi-squared test comparing the proportion of loss-of-function intolerant genes (defined as LOEUF < 0.35) among differentially expressed genes (DEGs) versus a background gene set. Results show that DEGs in large CNVs are significantly enriched for genes intolerant to loss-of-function variation, suggesting that transcriptional dysregulation in these CNVs disproportionately affects genes under strong functional constraint.*

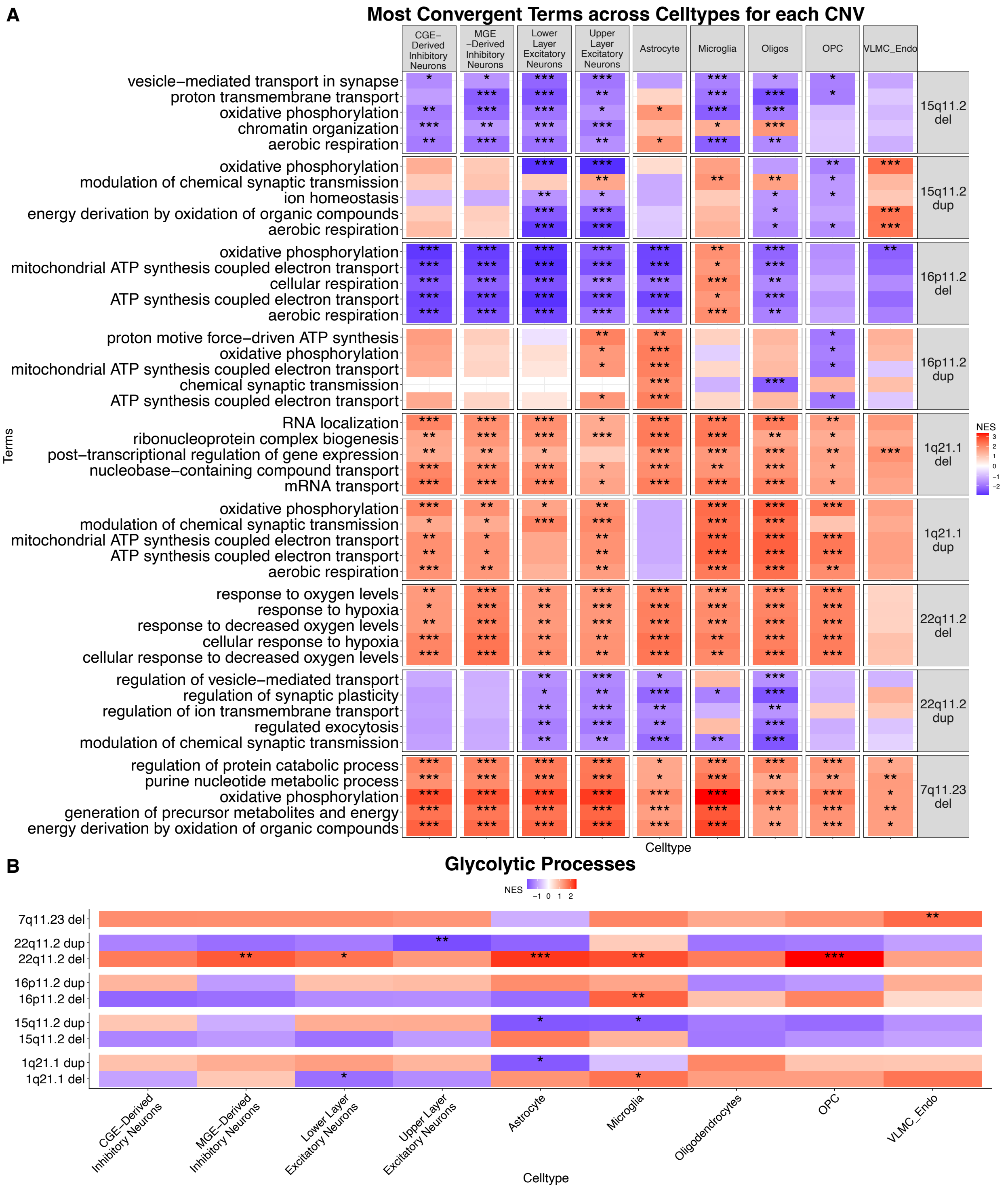


***Figure S11**: Convergent functional Enrichments for each CNV across cell types.* *(A) Heatmap of the top seven most frequently implicated term across the nine cell types for each CNV from GSEA[18,21]. Heatmap is colored by normalized enrichment scores (NES) with red indicating higher expression and blue indicating lower expression compared to non-carriers. FDR significance is marked with asterisks (FDR <0.05 = *, FDR <0.01 = **, FDR < 0.001 = ***). (B) Heatmap showing the Glycolytic Process (GO:0006096) GSEA results. The grid is colored by NES and annotated by FDR significance thresholds, as in A.*

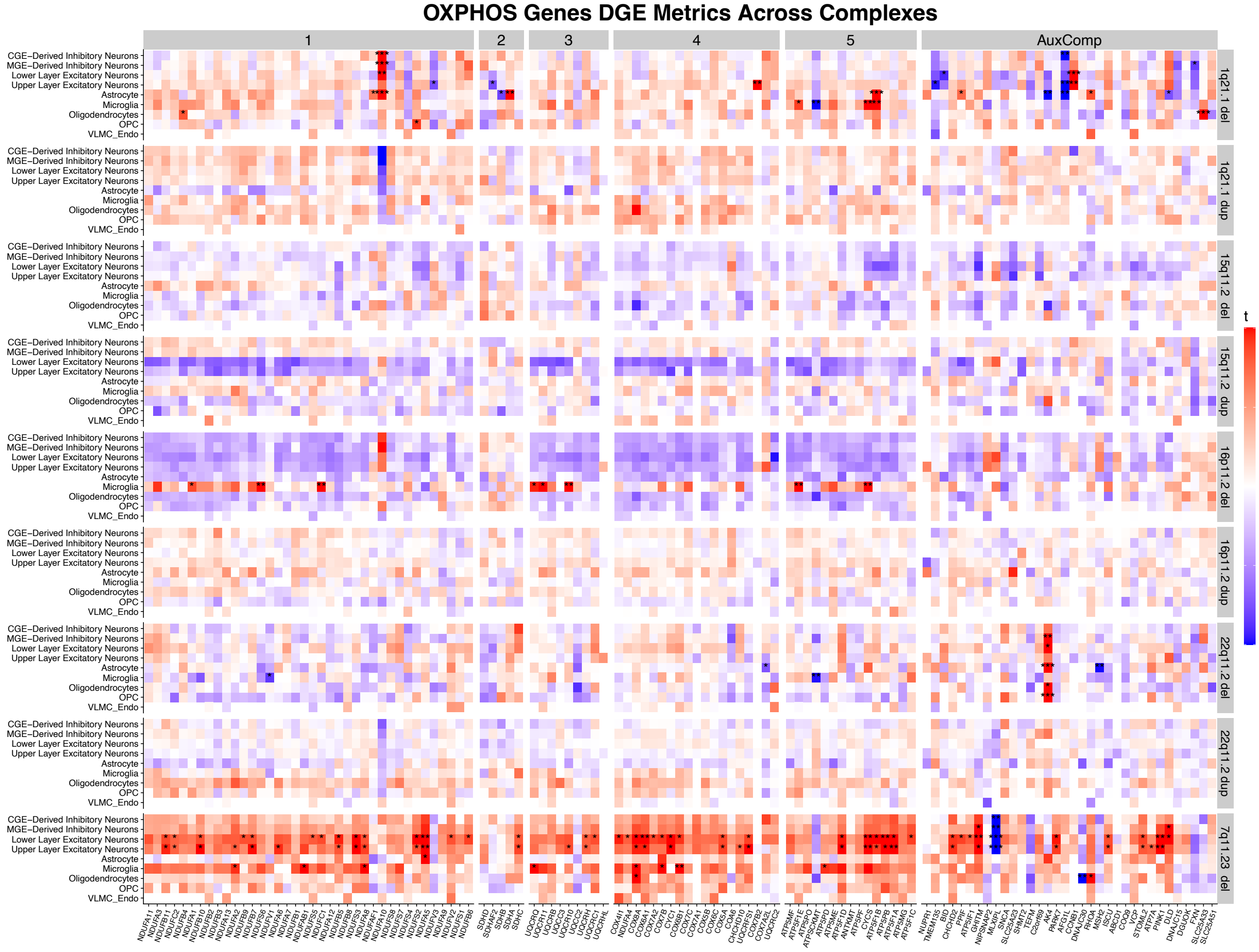


***Figure S12: Differential gene expression and functional enrichments of OXPHOS complexes.*** *(A) T-statistics for genes included in the OXPHOS term (GO:0006119), which were significant in one or more of the 81 models. Heatmap is separated by CNV (y-axis) and OXPHOS complexes (x-axis), as defined by the KEGG pathway [28]. Results from all 81 models are reported for all nine CNVs and cell types (y-axis). Genes within the auxiliary components, not specific to a complex, as the sixth column. FDR significance labeled with * (FDR <0.05 = *, FDR <0.01 = **, FDR < 0.001 = 0.0001). The 7q11.23 deletion shows upregulation of most genes inside the pathway, apart from the two genes in the 7q11.23 region, DNAJC30 and MLXIPL. For the 22q11.2 deletion, the most significant gene across cell types is AK4, which is known to be upregulated during chronic hypoxia [29].*

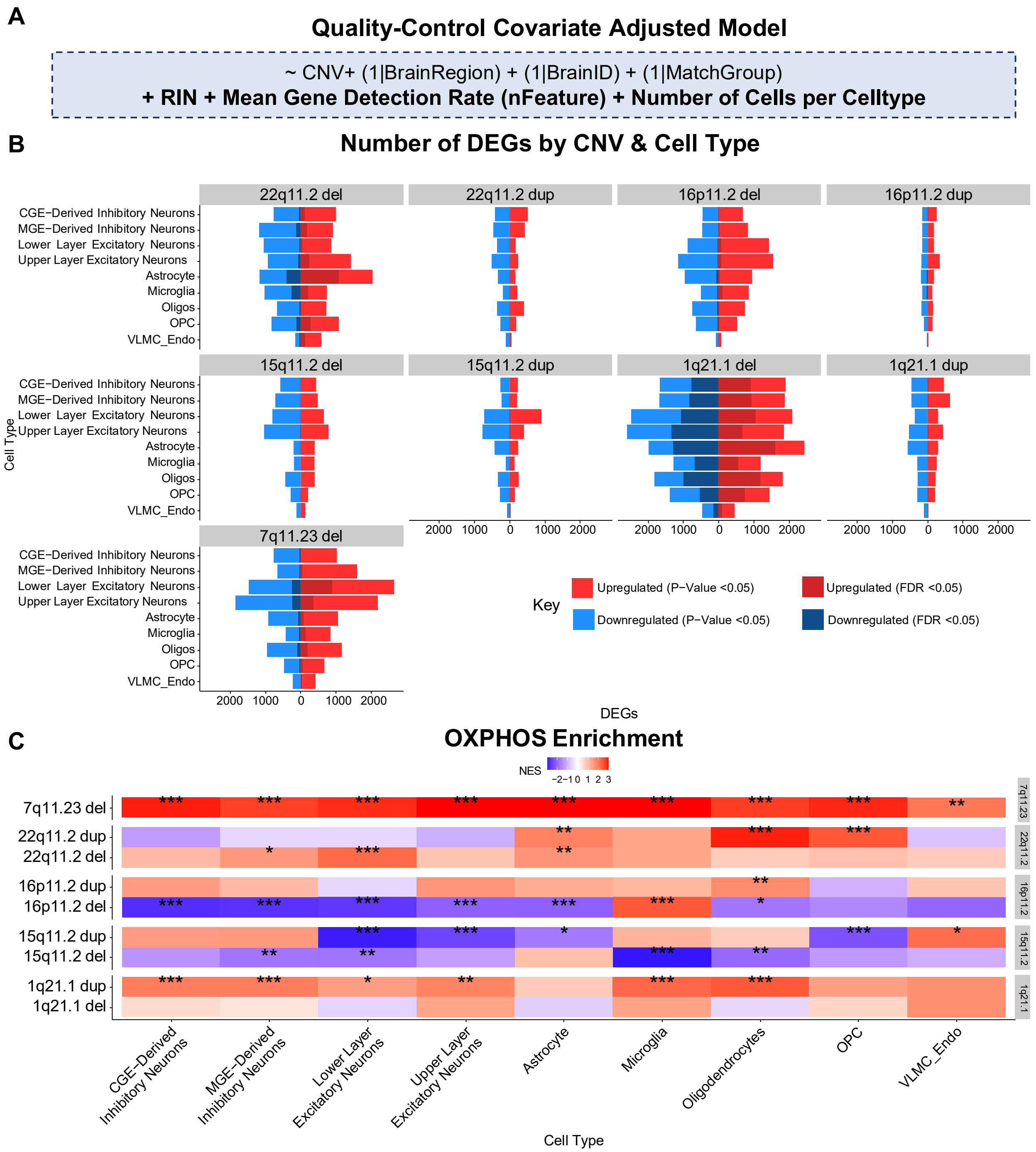


***Figure S13: QC-covariate adjusted model differential gene expression results.*** *(A) Outline of model: ~CNV + (1|BrainRegion) + (1|BrainNumber) + (1|MatchGroup) + RIN + nFeature + Cell.Type.Counts (B) Barplot of DEG counts at nominal and FDR-significance. (C) Heatmap of OXPHOS GSEA results across cell type (x-axis) and CNV (y-axis). Heatmap is colored by NES and asterisks show FDR significance thresholds * (FDR <0.05 = *, FDR <0.01 = **, FDR < 0.001 = 0.0001).*

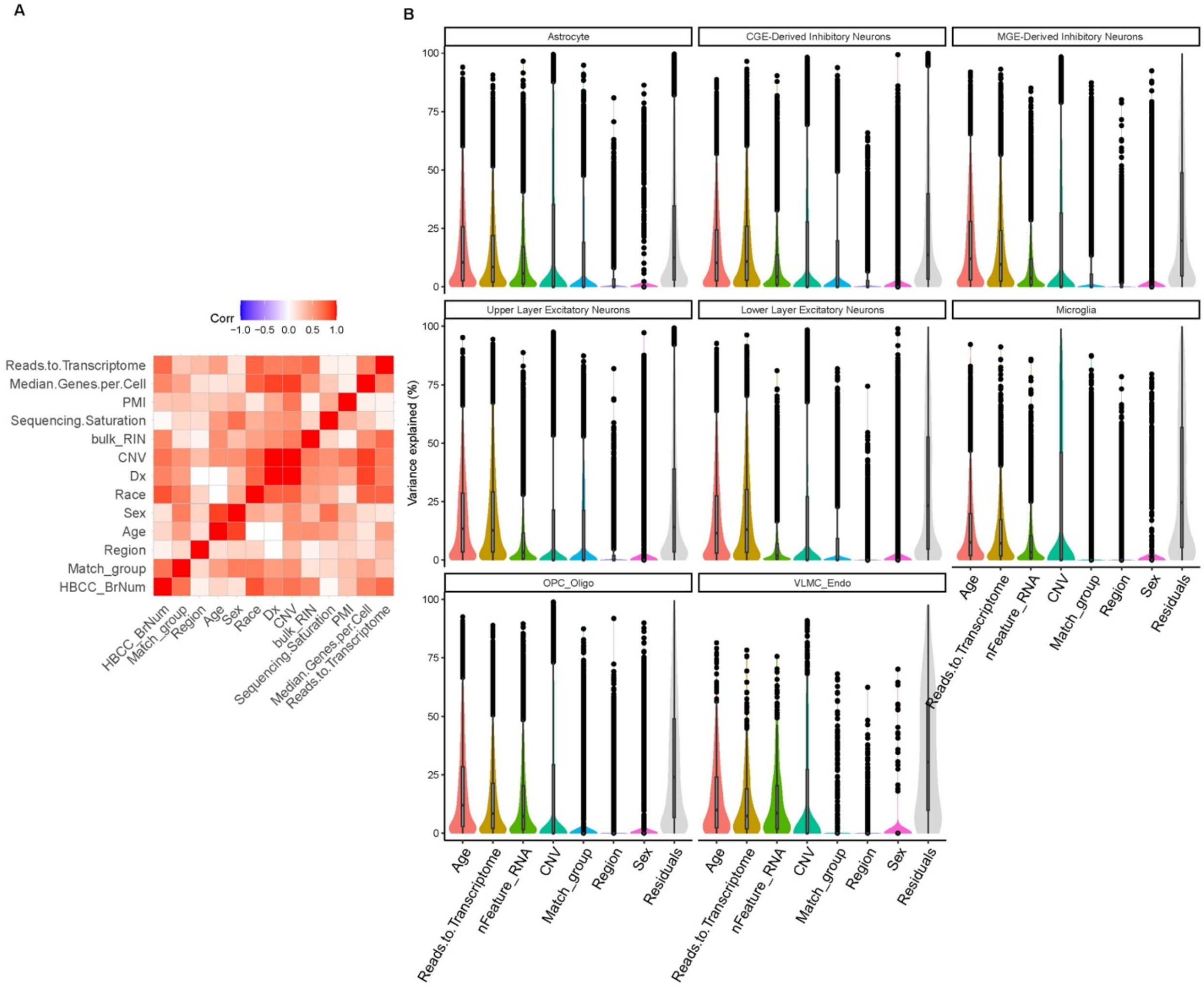


***Figure S14: Covariate selection for infant data.*** *(A) Correlation of covariates using canCorPairs from variance partitioning [27] (as in Figure S7). (B) Variance partitioning analysis showing the percentage of variance explained across genes for selected covariates (see also Table S3).*

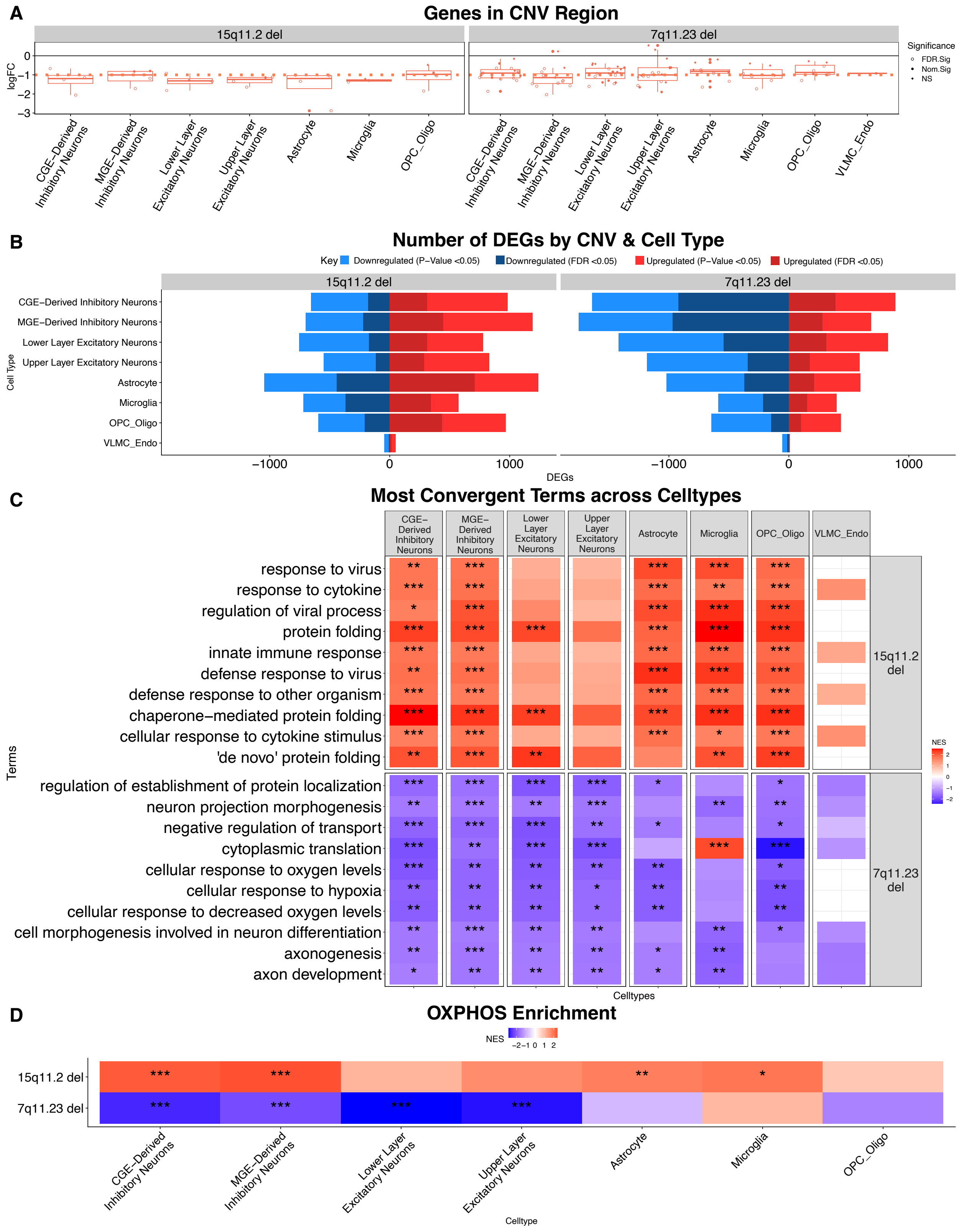


***Figure S15: Infant data differential gene expression and functional enrichment results.*** *(A) logFC of CNV genes from infant DGE models. CNV genes were obtained using same coordinates described in Figure 2 (see also Table S1, S2). CNV genes are downregulated (negative logFC) in both deletion*

*carriers, as expected. (B) Number of genes meeting nominal (p-value <0.05) and FDR significance (FDR <0.05). (C) GSEA[18] results of the top 5 most frequently implicated pathways enriched at FDR <0.05 across CNVs were selected for 15q11.2 and 7q11.23 deletion infant carriers across cell types (D) Heatmap showing the OXPHOS (GO:0006119) functional enrichment results, eight cell types (x-axis) and two CNVs (y-axis). The grid is colored by NES and annotated by FDR significance thresholds (FDR <0.05 = *, FDR <0.01 = **, FDR <0.001 = ***).*

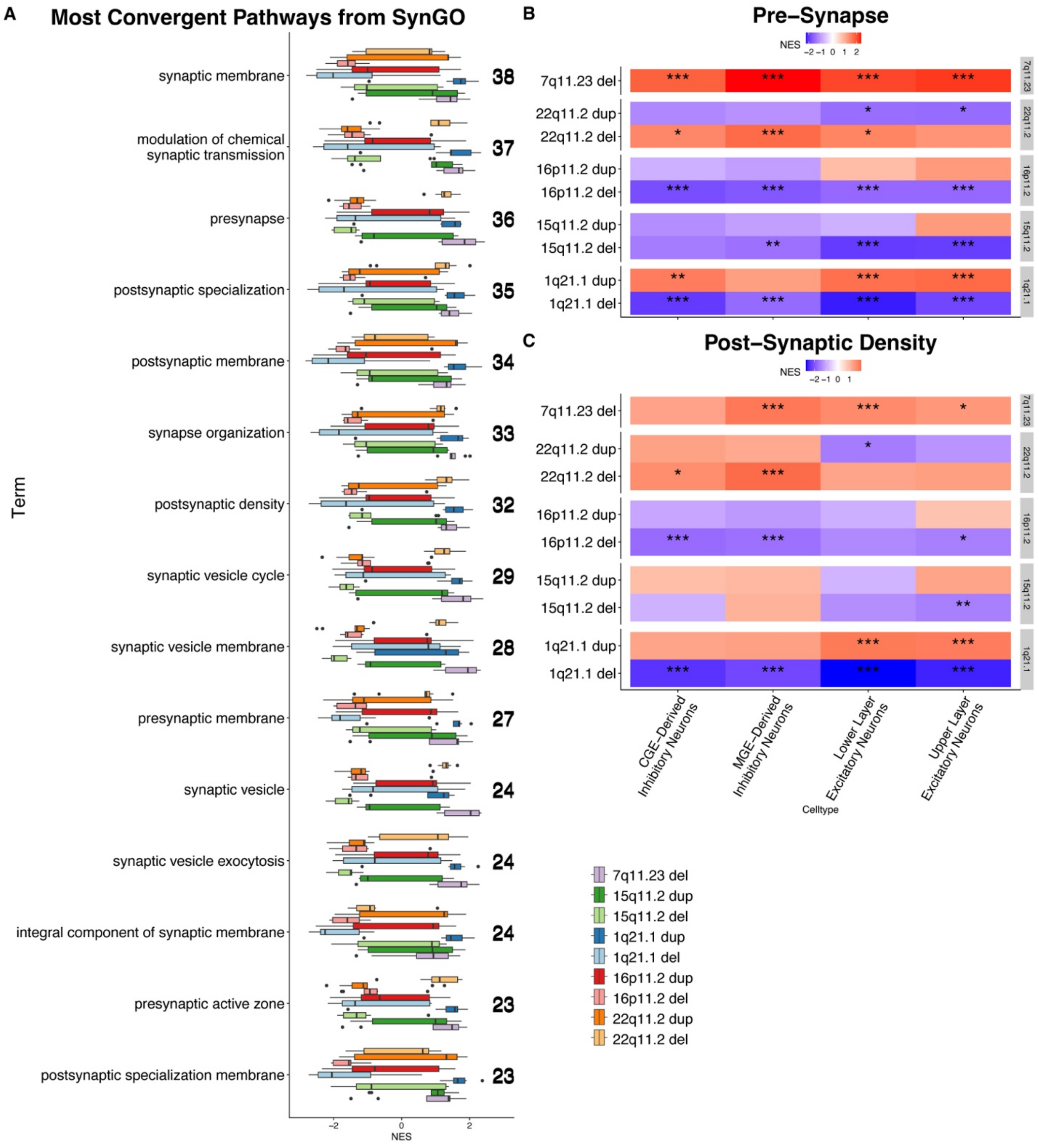


***Figure S16: Synapse-related terms from SynGO**. (A) Boxplot depicting NES (x-axis) of the top 15 most frequently implicated terms from the SynGO[25] database. Terms are ordered by frequency, with the number of models with FDR <0.05, shown on the right (of the total 81 models). (B) Heatmap showing the*

*GSEA results for presynapse (GO:0098793) results, across nine cell types (x-axis) and nine CNVs (y-axis). The grid is colored by NES and asterisk show FDR significance thresholds (FDR <0.05 = *, FDR <0.01 = **, FDR <0.001 = ***). The 22q11.2 deletion and duplication diverge strongly in direction for neurons (C) Heatmap showing the GSEA results for postsynaptic density (GO0014069)- colored and labelled as in B.*

### Supplementary Tables:

| CNV | Start | End |
|---|---|---|
| 1q21.1 | 147105904 | 147917509 |
| 7q11.23 | 73330452 | 74728172 |
| 15q11.2 | 22782170 | 23040134 |
| 16p11.2 | 29638676 | 30188531 |
| 22q11.2 | 18924718 | 21111383 |

***Table S1: Reference CNV coordinates derived from ClinGen[1].***

***Table S2: CNV genes for each CNV and cell type DGE model.***

***Table S3: Sample metadata overview.*** *Demographic covariate overview for adults and infants, including age, diagnosis, ethnicity, sex, etc.*

***Table S4: Sample metadata metrics from Cellranger.***

***Table S5: Results of linear mixed-effects model analysis for cell-type proportion testing.*** *This supplementary table contains nine individual sheets, each corresponding to a specific cell type. For each cell type, the results of the linear mixed-effects model are reported, along with the Shapiro-Wilk test p-value for normality and the transformation method applied to the cell-type proportions prior to modeling.*

***Table S6: Results of linear mixed-effects model analysis for cell-type count testing****. Similar to Table S5, this table provides linear mixed-effects model results, but here applied to cell-type counts. Each sheet includes the Shapiro-Wilk test p-value and details on the transformation used for count data.*

***Table S7: Proportion of genes, retained after filtering by expression, differentially expressed (FDR < 0.05) for each DGE model.*** *This table reports the percentage of genes that meet the false discovery rate (FDR) threshold of 0.05 following the application of filtering criteria in each DGE model.*

***Table S8: Differential gene expression results for each CNV and celltype.*** *These tables report the logFC, t-statistics, p-values, and FDR-adjusted p-values for all genes in the respective DGE models.*

***Figure S9: GSEA results for each CNV and celltype.*** *These tables report the normalized enrichment scores, p-values, and FDR-adjusted p-values for GO terms.*
